# *NORAD* prevents axonal degeneration by orchestrating KLC1-mediated SFPQ transport via liquid-liquid phase separation

**DOI:** 10.64898/2026.01.06.697853

**Authors:** Shuo Han, Huan Chen, Jiayan Liu, Yang Wang, Shaoxiang Chen, Panpan Sun, Jing Jiang, Rubin Zhao, Ruijun Chen, Xiaoxian Zhang, Fuyu Duan, Li Rong, Wenkai Yi, A. B Kinghorn, Jian Yan, Renhao Li, Junjie Xia, Zhongquan Qi, Gaël Ménasché, Kwok-On Lai, Cora Sau Wan Lai, Pengcheng Zhou, Jian-Dong Huang, Qizhou Lian

## Abstract

Neuronal function requires precise long-distance axonal transport mediated by molecular motors and RNA-binding proteins like SFPQ, though regulatory mechanisms remain poorly defined. We identify long non-coding RNA *NORAD* as a master regulator of this process through liquid-liquid phase separation (LLPS). While SFPQ and kinesin-1 mediate cargo delivery, the specific RNA coordinating their interaction was unknown. We demonstrate *NORAD* directly binds kinesin light chain 1 (KLC1) and promotes SFPQ condensation into dynamic LLPS droplets, enabling efficient transport. CRISPR-assisted mapping and functional assays show *NORAD* depletion disrupts granule dynamics, impairs neuroprotective mRNA localization, and induces axonal degeneration. *In vitro* reconstitution confirms *NORAD*-KLC1 synergy enhances SFPQ phase separation, and neuron-specific knockout mice exhibit motor deficits with reduced neuronal density. These findings establish the first evidence of lncRNA *NORAD*-mediated LLPS in axonal transport, revealing a new paradigm for RNA-guided neuronal maintenance.

## Introduction

Neuronal polarization establishes one of the most fundamental functional architectures in the nervous system, requiring precise long-distance transport of proteins along axons to synaptic sites. This process, known as axonal transport, depends on molecular motors traversing microtubule networks and RNA-binding proteins(RNP) coordinating cargo delivery^1,2^. Despite its critical role in maintaining neuronal function, the regulatory mechanisms governing this process, particularly the involvement of specific RNA molecules, remain incompletely understood. Among key players, Splicing Factor Proline- and Glutamine-Rich (SFPQ) has emerged as a ubiquitous RNA-binding protein essential for neuronal polarity^3–5^. It mediates intracellular transport and local translation of target mRNAs such as Bcl-w and Lmnb2, which are assembled into ribonucleoprotein (RNP) granules for delivery to distal axons^3,6–9^. Our previous study suggested that vesicles need to be transferred from microtubules to actin tracks to reach their destination by binding with motor proteins^10^. Recent studies established that SFPQ interaction with the kinesin-1 motor complex (KIF5A/KLC1) is required for axon survival^11^and is RNA-dependent^4^, as RNase treatment disrupts this interaction and impairs axonal integrity. However, the identity of the RNA species mediating this process has remained elusive.

In axons, the efficient transport of cargo, including RNP granules and presynaptic vesicles, occurs along the narrow pathways of neurons^23^. The high concentration of cytoskeletal filaments, organelles, and proteins, combined with the spatial constraints imposed by the narrow geometry of the axon, necessitates that cargo be flexible rather than stiff to maintain steady transport that can be hindered by physical crowding^24,25^. Axons of the pyramidal tract in humans have been reported to exhibit diameters ranging from as thin as 0.3 μm^26^ to as wide as 20 μm, with a predominant proportion of slender fibers (approximately 84%) measuring less than 2 μm in diameter^27^. In axons, the size and shape of SFPQ granules remain constant at approximately 1 µm in diameter, and the granules expand and elongate as they migrate to avoid axonal crowding^28^. We hypothesize that the transport of SFPQ in neuronal axons in a liquid state may facilitate the navigation of cargo in a crowded environment.

The long non-coding RNA *NORAD* (noncoding RNA activated by DNA damage) represents a promising candidate for regulating axonal transport. Highly abundant in mammalian brains and evolutionarily conserved^12–14^, *NORAD* exhibits predominantly cytoplasmic localization and declines markedly during aging and neurodegeneration. For isntstance , *Norad*^−/−^ mice exhibit neuronal pathologies in the dorsal root ganglia, brain stem, and cerebellum, accompanied by an overall reduction in neuronal density in the spinal cord^15^. In addition, *NORAD* has been shown to decline rapidly in the brain during aging in humans with neuronal degeneration^16^. Previous studies have shown that RNA-driven liquid-liquid phase separation (LLPS) is a key mechanism through which *NORAD* promotes the activity of Pumilio (PUM) proteins and maintains genomic stability in mammalian cells^14^. However, the role and mechanism of *NORAD* in neuronal transport processes remains unexplored.

In this study we address the role and mechanism of *NORAD* in axonal transport. Utilizing CRISPR-assisted detection of RNA-protein interactions (CARPID), we identified *NORAD* as a direct interactor of kinesin light chain 1 (KLC1). Through a combination of *in vitro* phase separation assays, live-cell imaging, and neuron-specific knockout models, we demonstrate that *NORAD* facilitates the assembly of SFPQ into dynamic liquid-liquid phase separation (LLPS) condensates, which are crucial for efficient cargo transport. Our findings uncover a previously unrecognized RNA-guided mechanism that maintains axonal integrity and offer new insights into the pathogenesis of neurodegenerative disorders.

## Results

### Depletion of *NORAD* results in axonal degeneration in human pluripotent stem cells-derived neurons

A mouse ortholog of *Norad* (*2900097C17Rik or Norad*), exhibiting 61% nucleotide identity with its human counterpart, is clearly identifiable on mouse chromosome 2. Like the human transcript, mouse *Norad* shows minimal protein-coding potential as assessed by PhyloCSF, a metric that discriminates between coding and noncoding sequences based on their evolutionary signatures^64^. To define the expression profile of *NORAD/Norad* across the tissues, we found that both human and mouse *NORAD/Norad* are ubiquitously expressed throughout the body, with highest abundance in brain **(Fig. 1a)**. Using RNA sequencing in human induced pluripotent stem cells (hiPSCs/PSC), derived human neural progenitor cells (hNPCs/NSC) and human neurons (Neuron)^32^, we demonstrated that *NORAD*’s expression levels are stable and gradually increase **(Fig. 1b)**, consistent with the results validated by qPCR **(Fig 1c)**. Furthermore, RNA fluorescent in situ hybridization (RNA FISH) in hESC, hNSC and hNeuron **(Fig 1d)** and subcellular fractionation in hESC-derived neuron **(Supplementary Fig. 1a)** demonstrated an almost exclusive cytoplasmic^12^ localization of *NORAD* in both axon and dedrites **(Supplementary Fig. 1b)**. These results show that a potential involvement of *NORAD* in the intricate processes underlying neuronal function.

**Fig. 1:**
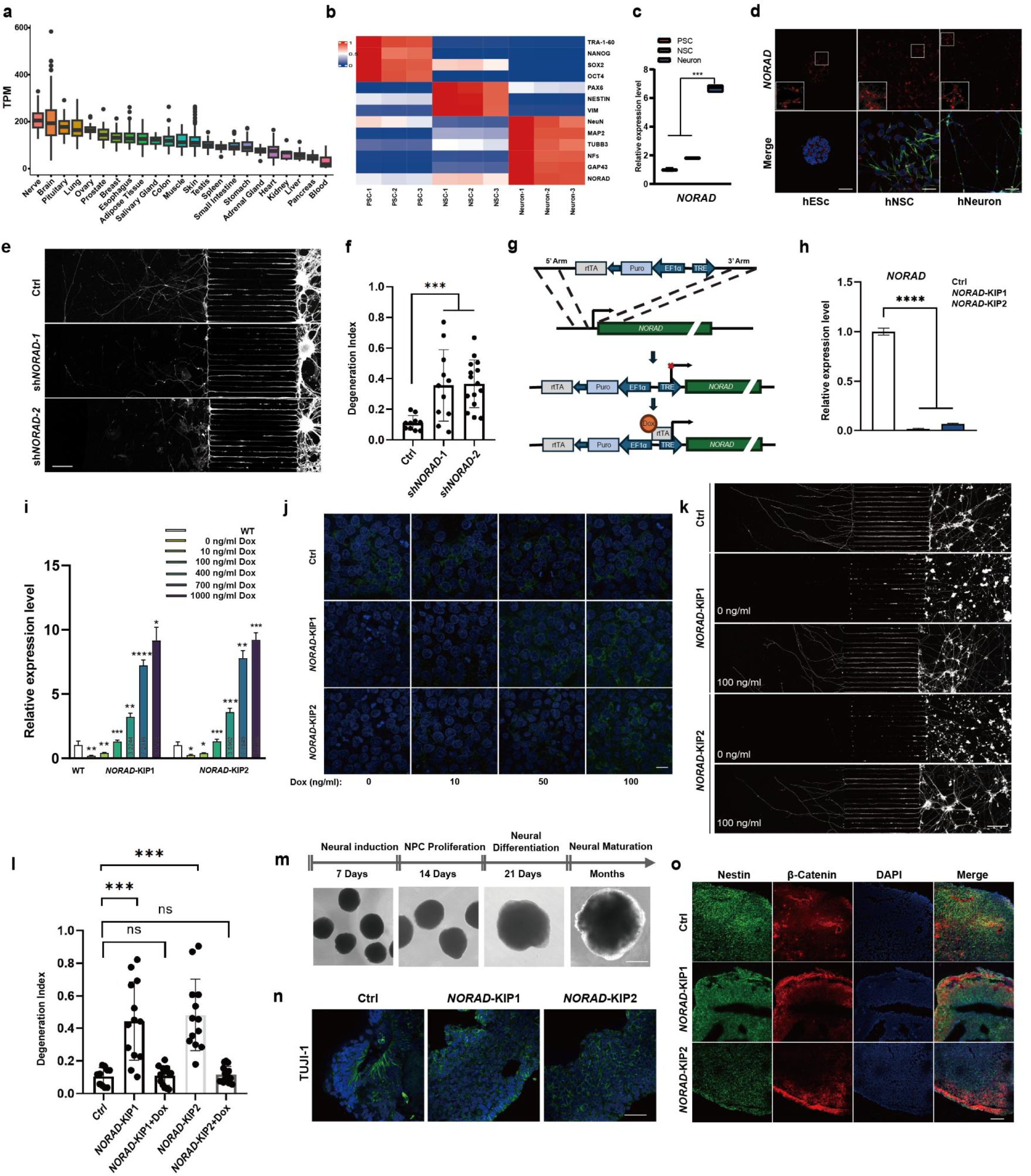
Depletion of *NORAD* results in axonal degeneration. **a** *NORAD* expression across 23 human tissues based on RNA-seq data from GTEx Consortium 65. **b** The RNA-seq (GSE115407) heatmap illustrates increasing *NORAD* expressions in PSC, NSC and Neuron. **c** qRT-PCR analysis of *NORAD* expression in PSC, NSC and Neuron (mean ± SEM, n = 3, biological replicates; \*\*\**p* < 0.001, unpaired t test). **d** Representative *NORAD* RNA FISH (red) images of hESC, hNSC (Nestin, green) and hNeuron (TUJI-1, green). Scale bar: 50 μm. **e** Staining of endogenous TUJI-1(White) in hESC-derived neurons grown in microfluidic chambers. Scale bar: 200 μm. **f** Quantification of axonal degeneration from K, which is calculated by using the ratio of the region of fragmented axons to total axon area degeneration index66. Control (Ctrl) (n = 10 captured images) or sh*NORAD*-1 (n = 11), sh*NORAD*-2 (n = 15) (mean ± SD; \*\*\**p* < 0.001, unpaired t test). **g** The inducible promoter cassette was knocked into the original promoter region of *NORAD* (*NORAD*-KIP). **h** qRT-PCR analysis of *NORAD* expression in *NORAD*-KIP1/2 hESCs (mean ± SEM, n = 3, biological replicates; \*\*\*\**p* < 0.0001, one-way ANOVA). **i** *NORAD* expression in *NORAD*-KIP1/2 hESCs with DOX treatment (mean ± SEM, n = 3, biological replicates; \**p* < 0.05, \*\**p* < 0.01, \*\*\**p* < 0.001, \*\*\*\**p* < 0.0001, one-way ANOVA). **j** Representative *NORAD* RNA FISH (green) images of hESC with Dox treatment. Scale bar: 20 μm. **k** Axons (TUJI-1, white) in Ctrl and *NORAD*-KIP1/2 neurons grown in microfluidic chambers treated with DOX. Scale bar: 200 μm. **l** Quantification of axonal degeneration from K, which is calculated by using the ratio of the region of fragmented axons to total axon area degeneration index66. Ctrl (n = 10 captured images) or *NORAD*-KIP1 (n = 13), *NORAD*-KIP1+DOX (n = 13), *NORAD*-KIP2 (n = 12), *NORAD*-KIP2+DOX (n = 13) (mean ± SEM; ns, not significant, \*\*\**p* < 0.001, one-way ANOVA). **m** Schematic of the protocol used to generate cortical organoids. Scale bar: 400 μm. **n** Representative immunostaining of Ctrl or *NORAD*-KIP1/2 brain organoids. Axon was identified with TUJI-1 (Green). Scale bar: 20 μm. **o** Representative immunostaining of Ctrl or *NORAD*-KIP1/2 brain organoids. Nestin (Green), β-Catenin (Red). Scale bar: 100 μm.

Previous reports have shown a significant decrease in *NORAD* expression levels with aging^65^. To investigate *NORAD*’s function in the nervous system, we generated small hairpin RNAs (shRNAs) against *NORAD* (sh*NORAD*-1 and sh*NORAD*-2) in an hESc-derived neuron **(Supplementary Fig. 1c)**. Importantly, knockdown of *NORAD* resulted in axon shortening in an hESC derived neuron **(Fig. 1e)**. The length of the axon was quantified by analyzing the ratio of the fragmented axons over the total axon occupied area, referred to as the degeneration index **(Fig. 1f)**, indicating that the level of *NORAD* expression was associated with axonal viability. We next performed a knock-in of an inducible promoter before the *NORAD* locus in hESC (NORAD-KIPs) to minimize the genetic influence of *NORAD* knockdown, replacing the endogenous promoter **(Fig. 1g)**. Subsequently, we selected two out of 100 monoclonal cell lines that were suitable for further experimentation **(Fig. 1h and Supplementary Fig. 1d)**. Genotyping results confirmed the successful insertion and separation of the inducible promoter from the endogenous promoter **(Supplementary Fig. 1d)**. In the absence of doxycycline (DOX), *NORAD* expression was reduced by at least 75% **(Fig. 1h)**, and its expression could be restored and overexpressed in a dose-dependent manner upon DOX addition **(Fig. 1i)**. RNA fluorescence in situ hybridization (RNA FISH) further confirmed that DOX could induce *NORAD* expression in hESC **(Fig. 1j)**.

Interestingly, *NORAD* knockout in hESC did not affect their pluripotency or karyotype **(Supplementary Fig. 1e-j)**. Knockdown of *NORAD* did not affect the morphology of hESC colonies or mRNA level of pluripotency genes **(Supplementary Fig. 1e-g)**. Additionally, cell cycle analysis revealed that genomic stability was not affected **(Supplementary Fig. 1h)**. This contrasts with previous reports that linked *NORAD* to genomic stability in tumor cells^12–15^. This discrepancy may be due to the enhanced capacity of ESC repair mechanisms to ensure maintenance of their genomic integrity^33^. Moreover, *NORAD*-KIP1/2 hESC-derived neural stem cells (NSCs) revealed no discernible effects on the potency in NSC induction **(Supplementary Fig. 1i)** or neural differentiation **(Supplementary Fig. 1j)**. Nonetheless, depletion of *NORAD* resulted in a neurodegenerative phenotype associated with impaired axonal length in *NORAD*-KIPs neurons without DOX after two-weeks of culture **(Supplementary Fig. 1k, l)**. Remarkably, axonal length was rescued by culturing with 100 ng/ml DOX, resulting in axon length comparable to that of the control (Ctrl) group neurons **(Fig. 1k, l)**, confirming that the expression level of *NORAD* was closely related to axonal growth.

To investigate *NORAD*’s impact further, we generated cerebral organoids to mimic complex brain circuits^30^ **(Fig. 1m)**. Immunostaining revealed that KO *NORAD* brain organoids exhibited incomplete structure and a noticeable reduction in size in the absence of DOX **(Fig. 1n)**. In the early stages of neuronal polarization, β-catenin is enriched at the axonal tip, contributing to local activation of the Wnt pathway that regulates axonal formation^34^. Nonetheless most β-catenin was located around the cell body and was associated with reduced axonal formation compared with the control cerebral organoids **(Fig. 1o)**. Altogether, these results indicate that defect in NORAD leads to neural degeneration.

### Identification of proteins interacting with *NORAD* by CARPID

To identify candidate proteins that directly interact with *NORAD* in living neurons, we employed CRISPR-assisted detection of RNA-protein interactions (CARPID) in hESC-derived neurons, followed by Mass Spectrometry (MS) **(Fig. 2a)**^35^. Therefore, we first designed two sets of gRNA arrays, each consisting of gRNA sequences separated by a 30-nucleotide direct repeat, targeting six adjacent loci on the *NORAD* transcript **(Supplementary Fig. 2a)** with validation of gRNA efficiency by Biotin RIP **(Supplementary Fig. 2b)**. A total of 1075 common targets were identified across three independent experiments, representing 84.9%, 82.5% and 85.3% of total hits in each individual test **(Fig. 2b)**. Fold changes from three biological replicates were quantified, with all significantly enriched proteins selected for further analysis **(Fig. 2c)**. These proteins were predominantly localized in the cytoplasm or in both cytoplasm and nucleus **(Fig. 2d)**, consistent with the observation of cytoplasmic localization of *NORAD* **(Fig. 1d and Supplementary Fig. 1a)**. Further localization analysis revealed a substantial proportion of cytoskeleton-related subcellular localizations among these groups **(Fig. 2d)**. KEGG analysis indicated frequent associations of these targets with neurodegenerative diseases **(Fig. 2e)**. All targets enriched in neurodegenerative diseases were illustrated, with cytoskeleton-localized kinesin components enriched in most neurodegenerative diseases, including Parkinson’s disease and Alzheimer’s disease. Among the cytoskeleton localized proteins, we observed that two components of kinesine-1, KLC1 and KIF5A **(Fig. 2f and Supplementary Fig. 2c)**, were associated with these neurodegenerative diseases and are also reported as key motor proteins that maintain neuronal axon outgrowth^4,11^.

**Fig. 2:**
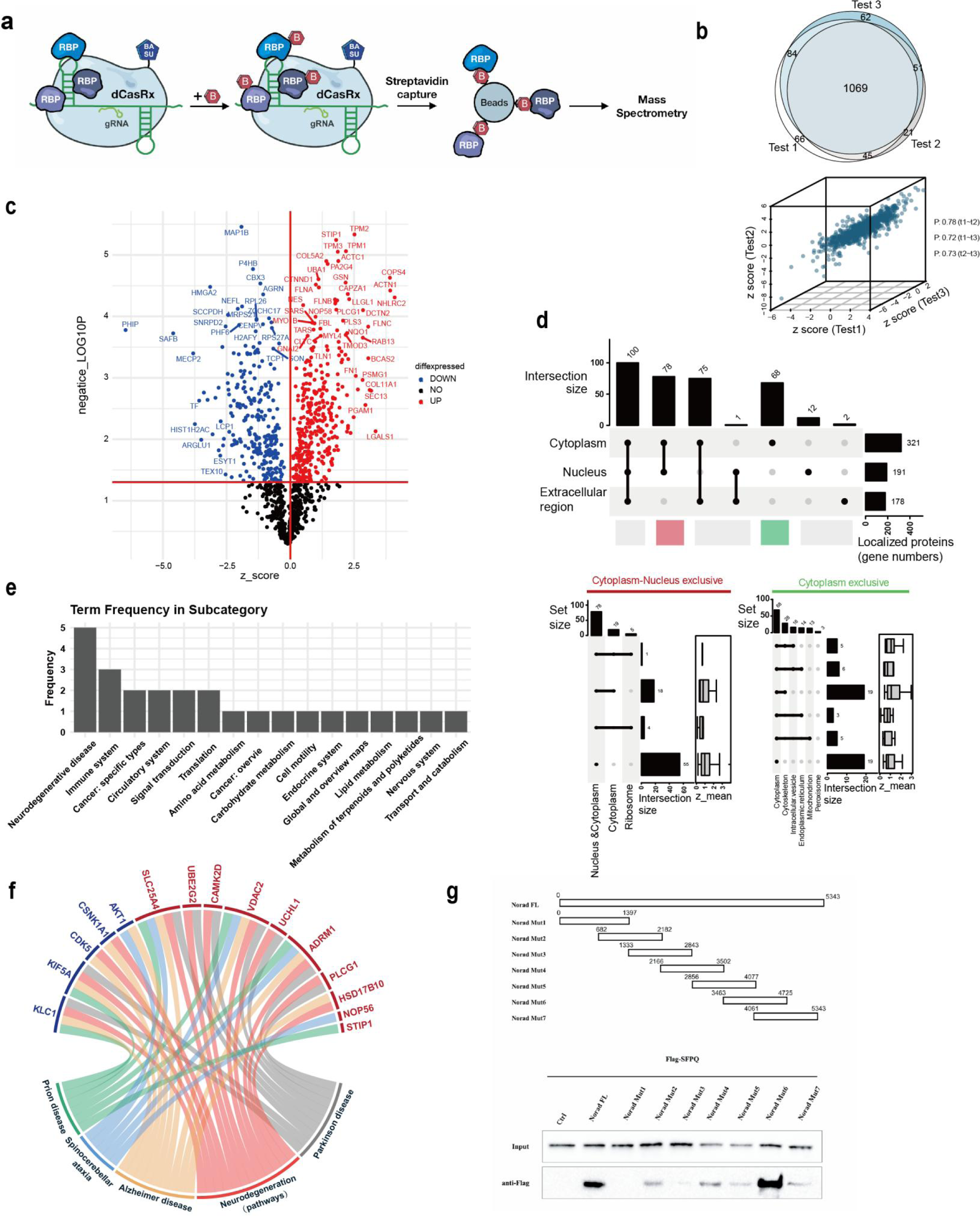
Identification of proteins interacting with *NORAD* by CARPID. **a** Schematic overview of CAPID done in hESC-derived neurons. Biotin is marked with ‘B’. **b** Reproducibility of detection of *NORAD*-interacting proteins by CARPID. The Venn diagram illustrates the overlap of *NORAD*-associated proteins with ≥ 2 peptides among the three independent experiments using the same gRNA set (Test1/Test2/Test3). The scatter plot of normalized log2-transformed label-free quantification intensities (Log2 LFQ) from three biological replicates demonstrated a reproducible result with high Pearson correlation coefficients (> 0.7 among either test). **c** The enrichment of *NORAD*-associated proteins. The volcano plot highlights the enriched *NORAD*-associated proteins (Red, p < 0.05, log2 LFQ > 0). **d** Significant cytoskeleton localization of enriched *NORAD*-interacting proteins. The subcellular localization analysis by SubcellularRVis is illustrated in UpSet plots. Upper: significant subcellular localization (p < 0.05) of all enriched *NORAD*-associated proteins. Lower: significant detailed subcellular localization (p < 0.05) of cytoplasm-nucleus exclusive and cytoplasm exclusive proteins sets. **e** The enriched neuron disease-associated KEGG terms of *NORAD*-interacting proteins. The bar chart illustrates the frequency of significantly enriched KEGG subcategory terms (p < 0.01). **f** The Chord diagram demonstrates enriched terms of neutron degenerative diseases and related genes from KEGG. **g** RNA pulldown assays were performed with truncated *NORAD* variants and followed by immunoblotting with the Flag antibody.

To explore the potential role of *NORAD* interactome in neurons, we analyzed high-confident interactions among all enriched cytoplasmic proteins. SFPQ was highlighted as a protein associated with KLC1 in the interaction network binding with *NORAD* **(Supplementary Fig. 2d, e)**^28^. To further analyse the interaction between *NORAD* and SFPQ, we constructed 7 biotinylated truncations of *NORAD* (1-1379nt, 682-2182 nt, 1333-2843nt , 2166-3502nt, 2856-4077nt, 3463-4725nt and 4061-5343nt) immunoprecipitated with purified Flag-SFPQ to define the *NORAD* and SFPQ binding. The Mut6 (3463-4725nt) truncation of *NORAD* exhibited the highest binding affinity for SFPQ compared with other truncations, as confirmed by the RNA pulldown assay **(Fig. 2g)**. SFPQ is predominantly located in the nucleus but also in the cytoplasm where it participates in distal axonal transport and in local translation^3^. Although SFPQ is less abundant in the cytoplasm, its significant role in neuronal axons suggests a potential interaction with *NORAD*, possibly forming a *NORAD*/SFPQ complex in axons.

### *NORAD* regulates the intracellular transport of SFPQ by binding with Kinesin- 1

As has been previously reported that the transport of SFPQ via its interaction with KLC1 is RNA-dependent^4,21^. To investigate whether *NORAD* regulates the interaction between Kinesin-1 and SFPQ, we used CRISPR-Cas13 to knockdown *NORAD* **(Supplementary Fig. 3a)**. This was followed by Kinesin-1 co-immunoprecipitation (CO-IP). Knockdown of *NORAD* resulted in dissociation of the SFPQ and Kinesin-1 interaction **(Fig. 3a)**, with a reduction in motor-cargo interactions similar to that observed with RNase treatment **(Supplementary Fig. 3b, c)**.

**Fig. 3:**
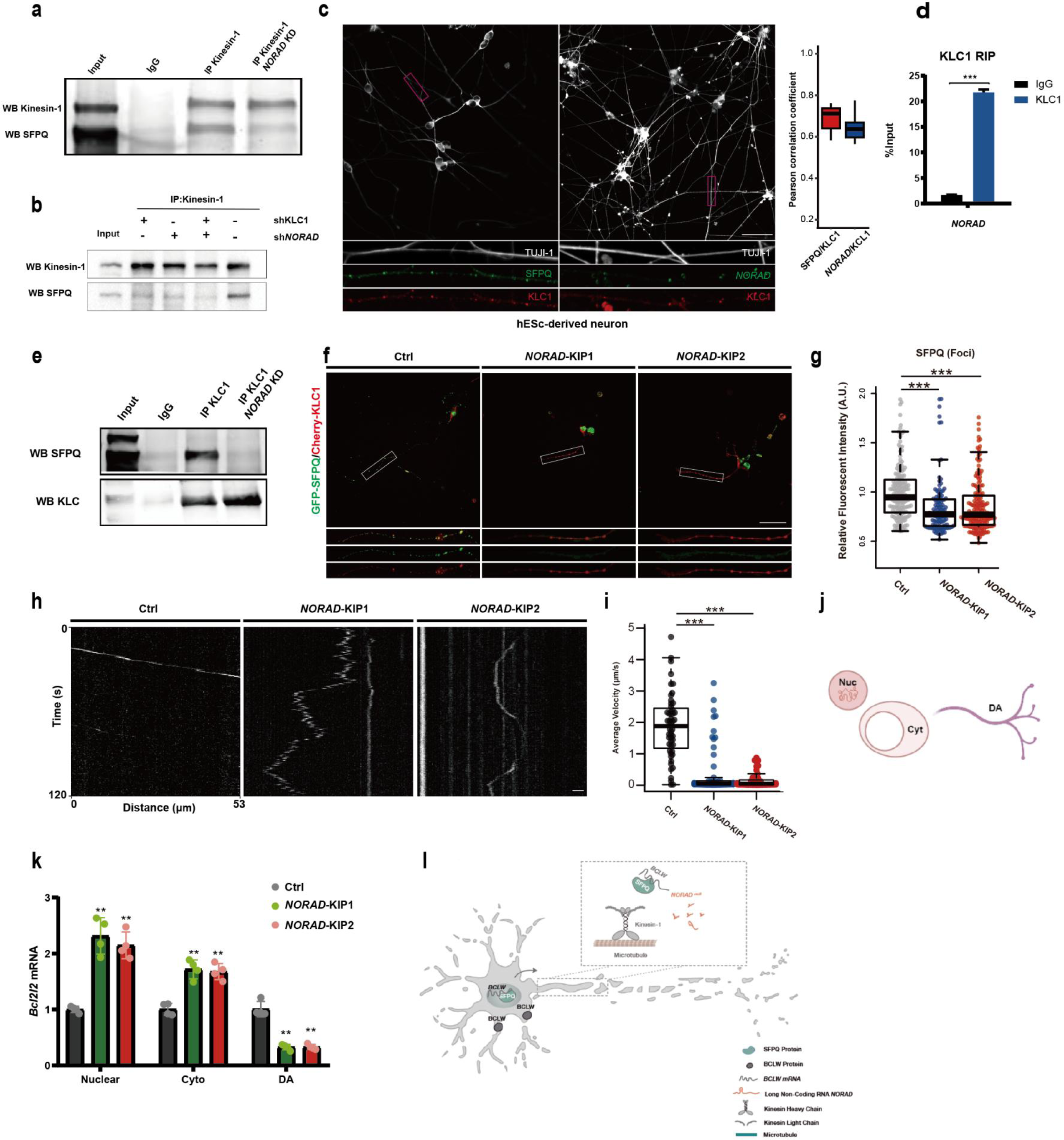
N*O*RAD regulates the intracellular transport of SFPQ by binding with KLC1. **a** Co-IP experiments were performed using Kinesin-1 antibody in *NORAD* knockdown cell. **b** Co-IP experiments were performed using Kinesin-1 antibody in KLC1 and/or *NORAD* knockdown cell. **c** Representative images of *NORAD* RNA FISH and immunostaining for SFPQ and KLC1 in hESC-derived neurons. TUJI-1 (white), KLC1 (red), SFPQ (green). The boxes outline regions shown for the higher magnification view of axons. Scale bar: 50 μm. **d** RIP analysis for endogenous KLC1 and *NORAD* in hESc-derived neuron. (mean ± SEM, n = 3, biological replicates, \*\*\**p* < 0.001, one-way ANOVA). **e** Co-IP experiments were performed using KLC1 antibody followed by *NORAD* knockdown. **f** Representative GFP-SFPQ (green) and Cherry-KLC1 (red) images in Ctrl or *NORAD*-KIP1/2 neurons. White boxes outline regions shown for the higher magnification view of axons. Scale bar: 50 μm. **g** Quantification of mean fluorescent intensities of SFPQ foci. Data points indicate individual foci of SFPQ fluorescence. (mean ± SEM, n = 3, biological replicates, \*\*\**p* < 0.001, one-way ANOVA). **h** Kymograph depicting GFP-SFPQ in Ctrl and *NORAD*-KIP1/2 derived neurons. Scale bars: 5 μm. **i** Average velocities of SFPQ droplets movement. Box-and-whisker plots show the average velocities of individual SFPQ droplets in WT and *NORAD*-KIP1/2 cells. (mean ± SEM, n = 3, biological replicates, \*\*\**p* < 0.001, one-way ANOVA). **j** Subcellular fractionation of mRNA from nuclei (Nuc), cytoplasm (Cyt) and distal axons (DA) from Ctrl or *NORAD*- KIP1/2 neurons grown in compartmentalized cultures. **k** qRT-PCR analysis of mRNA levels from Nuc, Cyt and DA from Ctrl or *NORAD*-KIP1/2 neurons. (mean ± SD, n = 4, biological replicates; \*\**p* < 0.01, unpaired t test). **l** The model of *NORAD* regulated the transport of SFPQ in neuron.

To verify whether Kinesin-1 regulated the transport of SFPQ through a cooperative interaction with KLC1 and *NORAD*, we conducted Kinesin-1 CO-IP assays following the knockdown of KLC1 or *NORAD* **(Supplementary Fig. 3d, a)**. Depletion of either KLC1 or *NORAD* attenuated the interaction between Kinesin-1 and SFPQ. A further reduction in binding affinity was observed when both KLC1 and *NORAD* were knocked down simultaneously **(Fig. 3b)**. These findings suggest that KLC1 and *NORAD* function as a protein-RNA complex, with *NORAD* serving as an adaptor to mediate and facilitate the transport of SFPQ.

To further investigate the potential synergistic effects of KLC1 and *NORAD* in axonal transport, we performed *NORAD* RNA-FISH along with immunostaining for SFPQ, KLC1, and TUJI-1 in hESC-derived neurons.The results showed that both SFPQ and *NORAD* were co-localized with KLC1 in axons and dendrites, with average Pearson correlation coefficients exceeding 0.6 **(Fig. 3c and Supplementary Fig. 3e)**. To determine whether KLC1 possessed RNA-binding capabilities, we conducted a KLC1 RNA immunoprecipitation (RIP) assay and demonstrated significant enrichment of *NORAD* by KLC1 **(Fig. 3d)**. Additionally, KLC1 CO-IP was performed with or without RNase treatment, followed by MS to evaluate the binding affinity between KLC1 and its cargo. This analysis revealed that SFPQ was significantly decreased among a total of 433 proteins, with 330 of them being significantly downregulated (LogFC2 < -1, p < 0.05) **(Supplementary Fig. 3f)**. These results suggest that KLC1 coordinates the transport of numerous potential cargoes through synergistic interaction with RNAs.

To further substantiate the hypothesis that the interaction between KLC1 and SFPQ is contingent upon the presence of *NORAD*, we conducted CO-IP experiments targeting KLC1. These experiments were meticulously carried out following the strategic knockdown of *NORAD* in neurons to ascertain the dependency of KLC1- SFPQ binding on *NORAD* expression. As a result, the binding affinity between SFPQ and KLC1 was significantly reduced, providing compelling evidence that *NORAD* is indispensable for this interaction **(Fig. 3e)**. To determine whether *NORAD* regulated SFPQ transport in axons, we introduced GFP-SFPQ and Cherry- KLC1 into *NORAD*-KIP 1/2 hESCs **(Supplementary Fig. 3h, i)** and subsequently differentiated these cells into neurons. This experimental setup enabled us to track the dynamics of SFPQ and KLC1 in the presence and absence of *NORAD*, thereby elucidating the role of *NORAD* in the axonal transport of SFPQ. In both *NORAD*- KIP1 and *NORAD*-KIP2 neurons, most of the SFPQ was localized within the neuronal cell body rather than the distal axon compared with the control group **(Fig. 3f)**, even though depletion of *NORAD* did not reduce the protein expression of KLC1 or SFPQ **(Supplementary Fig. 3g)**. This suggests that *NORAD* plays a crucial role in the proper localization and transport of SFPQ within the axon. The quantified fluorescent intensities of SFPQ and KLC1 axonal foci significantly decreased in *NORAD*-KIP1 and *NORAD*-KIP2 cells **(Fig. 3g)**, indicating that the transport of SFPQ was disrupted in *NORAD*-KIP cells. This result is consistent with the phenotype observed in KLC1 knockdown neurons **(Supplementary Fig. 3j)**^11,36–39^.

To validate the role of *NORAD* in mediating the transport of SFPQ, we performed live cell imaging to track SFPQ granule movement along distal axons. Kymograph of SFPQ granules in wild-type cells displayed relatively uniform and rapid movement trajectories, while *NORAD*-KIP cells exhibited frequent back-and-forth and rotational movements in the axon **(Fig. 3h)**. The average velocity of SFPQ granules was significantly decreased in *NORAD*-KIP cells **(Fig. 3i).** This aberrant movement pattern in *NORAD*-KIP cells further supports the notion that *NORAD* is crucial for the efficient and directed transport of SFPQ granules along axons.

SFPQ is essential for the coassembly and axonal trafficking of *Bcl2l2 (B-cell lymphoma 2-like 2)* mRNAs into RNA granules, and a lack of *Bcl2l2* results in a distinct pattern of axonal degeneration^40,41^. To determine whether *NORAD* regulates SFPQ-mediated mRNA transport in axons, we performed subcellular fractionation of mRNAs from the nuclei, cytoplasm, and distal axons **(Fig. 3j)**. *NORAD* depletion led to the accumulation of *Bcl2l2* mRNA in the nuclear fraction **(Fig. 3k)**, suggesting that *NORAD* is essential for axonal transport of *Bcl2l2* mRNA. Together, these results suggest that *NORAD* plays a role in mediating the transport of SFPQ and is crucial for the maintenance of neurons **(Fig. 3l)**.

### *NORAD* facilitates axonal transport of SFPQ via liquid-liquid phase separation

In axonal transport, the high concentration of cytoskeletal filaments, organelles, and proteins, combined with the spatial constraints imposed by the narrow geometry of the axon, can result in stalling of cargoes similar in diameter to the neuronal processes. This stalling occurs due to the varying axon diameters that range from as thin as 0.3 μm to as wide as 20 μm^26,27,42–45^. The diameter of SFPQ granules is approximately 1-2 μm^4^ so may cause physical congestion of trafficking within neuron axons. Previous studies have demonstrated that the ability of SFPQ to suppress TGF-β responses is dependent on its prion-like domain (PrLD) that facilitates liquid-liquid phase separation (LLPS)^46,47^. We hypothesized that SFPQ may be conveyed as liquid condensate during axonal transport.

To investigate whether the SFPQ protein undergoes phase separation in neuronal axons, we examined the fusion and coalescence of GFP-SFPQ during axonal transport. Our observations demonstrated that GFP-SFPQ exhibits a liquid-like state in the dense phase, characterized by its ability to undergo fusion **(Fig. 4a)**. Additionally, GFP-SFPQ foci exhibited liquid-like properties, including rapid exchange with the surrounding area, as demonstrated by fluorescence recovery after photobleaching (FRAP) with an average recovery half-time of 10 seconds **(Fig. 4b**). These results indicate that the process of SFPQ axonal transport occurs in a liquid-liquid phase separation manner.

**Fig. 4:**
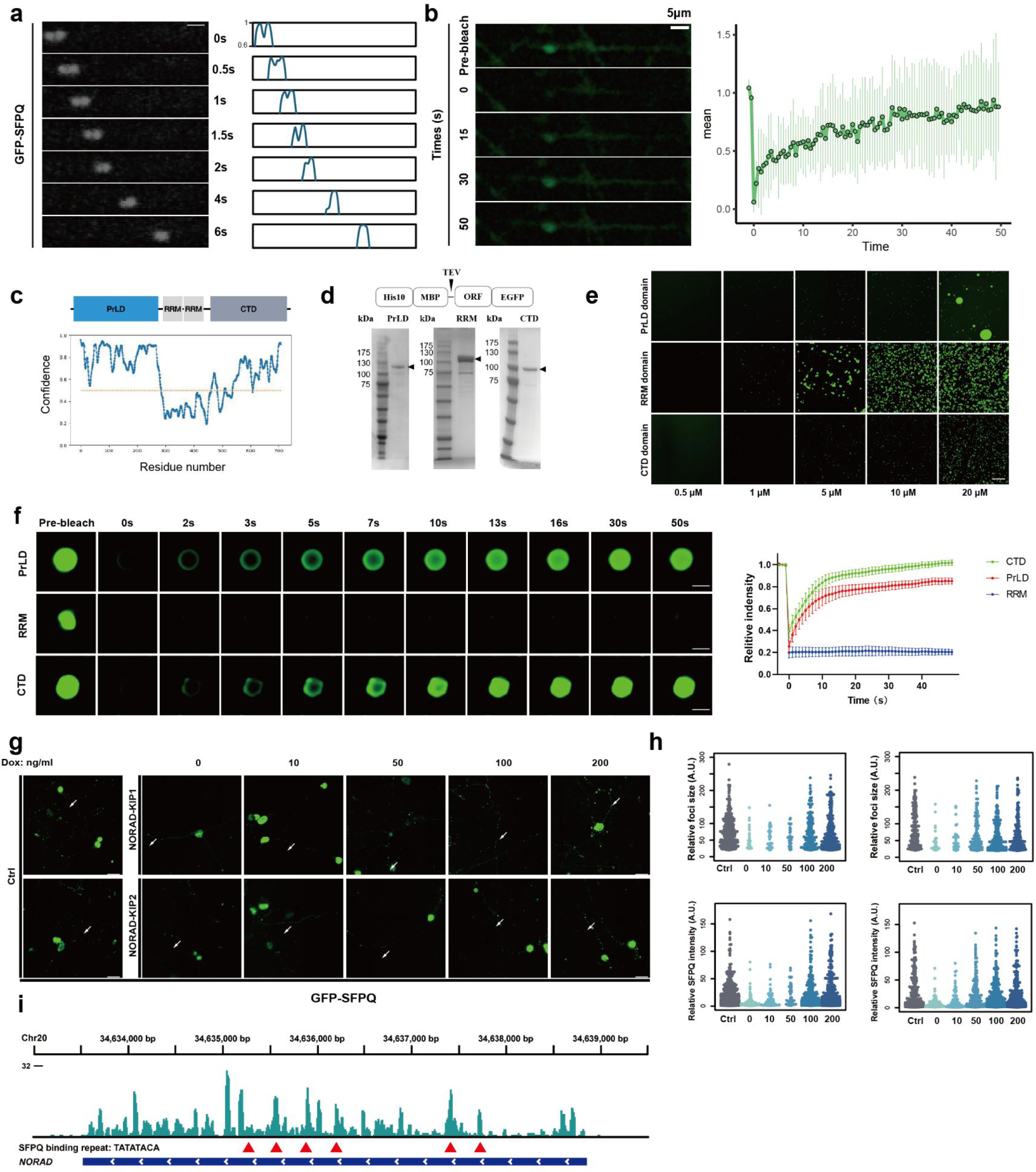
N*O*RAD facilitates axonal transport of SFPQ via liquid liquid phase separation. **a** Time lapse images of GFP-SFPQ are motility and undergo fusion in hESC-derived neural axon. Scale bar: 5 μm. Quantified and normalized fluorescence peak intensities are illustrated. **b** Images (left) and quantification (right) of GFP-SFPQ FRAP. Fluorescence intensities plotted relative to pre-bleach time point (t = −1 s). Data shown as mean ± SD (n = 9 droplets). Scale bar: 5 μm. **c** Predicted SFPQ domains contribute to LLPS. **d** Proteins purifications of PrLD domain, RRM domain, and CTD domain. Validated by SDS-PAGE. **e** PrLD domain, RRM domain and CTD domain form droplet in different concentrations. Scale bars: 20 μm. **f** Confocal images (left) and quantification (right) of PrLD domian, RRM domain and CTD domain of SFPQ droplet FRAP (1 μM). Fluorescence intensities plotted relative to pre-bleach time point (t = -5 s). Data shown as mean ± SD (n = 3 droplets). Scale bar: 5 μm. **g** Confocal images of GFP-SFPQ droplet in Ctrl or *NORAD*-KIP1/2 neural axons. Scale bars: 20 μm. **h** Quantification of relative sizes and intensities of SFPQ condensates with increased *NORAD* expression. Box-and-whisker plots show the relative foci sizes and intensities of SFPQ (****p < 0.0001). **i** Schematic of *NORAD* transcripts. Mammalian sequence conservation shown in green (UCSC Genome Browser hg38 PhastCons track). Locations of repeats indicated with red arrowheads.

To confirm the LLPS formation of SFPQ, we conducted *in vitro* assays using purified proteins. SFPQ contains a prion-like domain (PrLD), a coiled-coil RNA recognition motif (RRM), and intrinsically disordered regions in the C-terminus domain (CTD) **(Fig. 4c)**. We expressed and purified EGFP-tagged SFPQ domains fused to TEV protease-cleavable maltose binding protein (MBP) and tandem histidine tags **(Fig. 4d)**. After TEV cleavage of the MBP and His tags, the PrLD, RRM and CTD domains of SFPQ formed liquid-like droplets when incubated at a high concentration (5μM) with polyethylene glycol (PEG), a molecular crowding reagent **(Fig. 4e)**. These droplets exhibited rapid exchange with the bulk phase, as evidenced by their rapid recovery in fluorescence after photobleaching (FRAP) tests **(Fig. 4f)**. This indicates the liquid-liquid phase separation properties of SFPQ and suggests a mechanism in the liquid state for its transport within neuronal axons.

Moreover, by expressing GFP-SFPQ in *NORAD*-KIP neurons, SFPQ was recruited into droplets in a manner that was dependent on the level of *NORAD,* which expression is controlled by DOX **(Fig. 4g)**. GFP-SFPQ condensates were quantified in terms of foci size and fluorescent intensity, and were significantly decreased in *NORAD*-KIP cells and rescued by inducing *NORAD* re-expression **(Fig. 4h)**. These findings indicate that *NORAD* regulates SFPQ condensate formation in axons.

To identify the binding sites of SFPQ on *NORAD*, we utilized SpyTag-based crosslinking and immunoprecipitation (SPY-CLIP) with human SPY-tagged SFPQ^48^ to detect specific SFPQ binding interactions in neurons **(Supplementary Fig. 4a-c)**. The results showed that SFPQ binds to various types of RNAs, including 20% lncRNAs with multiple binding sites **(Supplementary Fig. 4d)**. We characterized the SFPQ binding sequence on *NORAD* using Spy-Clip **(Fig. 4i)** and found that the binding sequences (TATATACA) aligned well with previous findings of PUM- *NORAD* binding sites^12,14,49^. Together, our result shows that *NORAD* facilitates the partitioning of SFPQ into liquid-like droplets in axonal transport.

### *NORAD* collaborates with Kinesin-1 to promote the LLPS of SFPQ-PrLD domain

To determine whether the binding domains of SFPQ were promoted by *NORAD*, we expressed and purified various SFPQ fragments *in vitro* **(Fig. 4d).** In the absence of PEG-3350, only the SFPQ PrLD domain formed liquid-like droplets under the concentration conditions tested, and these condensates were promoted by *NORAD* **(Fig. 5a)**. We also purified the ΔPrLD domain **(Supplementary Fig. 5a)** and observed that the fluorescence of the condensates did not recover following fluorescence recovery after FRAP **(Fig. 5b)**. Additionally, the droplet could not be promoted by *NORAD* in the absence of the PrLD domain **(Fig. 5c)**. These results demonstrate that the LLPS of SFPQ is promoted by *NORAD* through formation of the PrLD domain. Furthermore, we labeled Cy3-*NORAD* and co-cultured it with GFP-PrLD SFPQ (10 μM) and revealed that the LLPS of the PrLD domain, in contrast to the RRM or CTD domains, was progressively enhanced with increasing concentrations of *NORAD* (from 0.05 to 10 nM) **(Fig. 5d and Supplementary Fig. 5b)**.

**Fig. 5:**
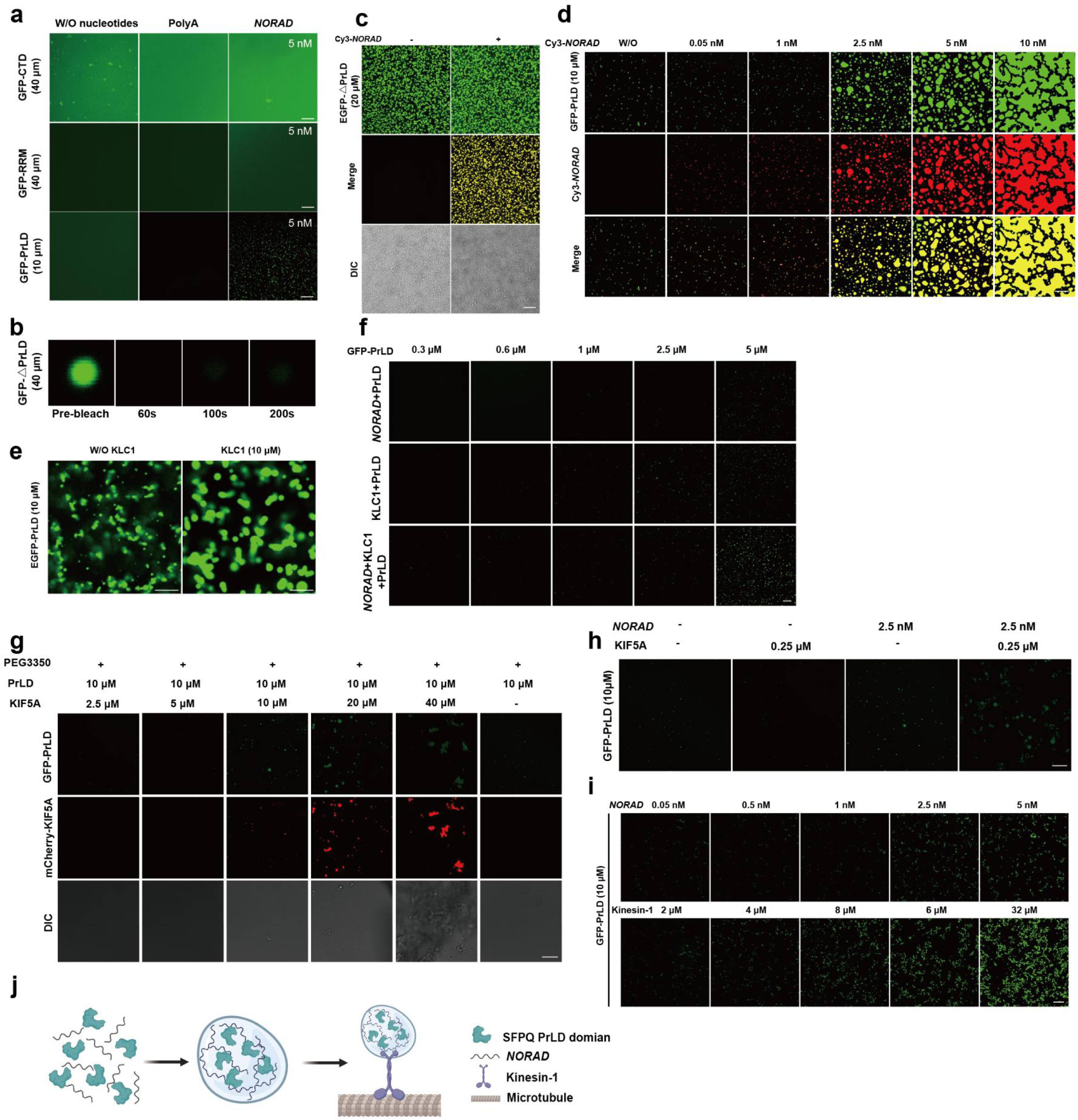
N*O*RAD collaborates with Kinesin-1 to promote the LLPS of SFPQ-PrLD domain. **a** Confocal images of GFP-tagged CTD, RRM, and PrLD domain droplets, with and without *NORAD*, in the absence of PEG-3350. Scale bars: 20 μm. **b** Images of GFP-◻PrLD domain of SFPQ FRAP. Fluorescence intensities plotted relative to pre-bleach time point (t = -5 s). **c** Confocal images of GFP-◻PrLD domain droplet with or without *NORAD*. Scale bars: 20 μm. **d** Confocal images of GFP-PrLD (green) droplets with different concentrations of Cy3-*NORAD* (red). Scale bars: 20 μm. **e** Confocal images of 10 μM EGFP-PrLD (green) droplets with or without recombinant 10 μM KLC1. Scale bars: 20 μm. **f** Confocal images of GFP-PrLD (green) droplets at varying concentrations in the presence of recombinant KLC1 and/or *NORAD*. Scale bars: 20 μm. **g** Confocal images of 10 μM GFP-PrLD (green) droplets with different concentrations of mCherry-KIF5A (red). Scale bars: 50 μm. **h** Confocal images of 10 μM GFP-◻PrLD (green) droplets with or without *NORAD* or KIF5A. Scale bars: 20 μm. **i** Confocal images of 10 μM GFP-PrLD (green) droplets with different concentrations of Kinesin-1. Scale bars: 20 μm. **j** Summary of the *NORAD* mediated axonal transport model.

There are multiple kinesin heavy chain isotypes and kinesin light chain isotypes in axons^50^. Given that KLC1 is involved in various cellular functions, including the transport of RNA, RNA granules, and proteins^21,51–55^, we propose that the function of KLC1 extends beyond RNA transport and may potentially collaborate with RNA to facilitate the LLPS of certain cargoes, thereby enhancing the efficiency of axonal transport. We initially purified Flag-tagged KLC1 protein **(Supplementary Fig. 5c)** and determined that a concentration of 10 µM KLC1 was sufficient to promote the phase separation of SFPQ PrLD **(Fig. 5e)**. Importantly, the simultaneous addition of KLC1 and *NORAD* significantly enhanced the phase separation of SFPQ PrLD compared with treatment with either KLC1 or *NORAD* alone **(Fig. 5f)**.

Previous studies have demonstrated that the transport of SFPQ-RNA granules by KIF5A/KLC1 motors is crucial for axon survival ^4^. Consistently, we also found that *NORAD* selectively bound to both KIF5A and KLC1 **(Supplementary Fig. 2d)**. We purified mCherry-tagged KIF5A **(Supplementary Fig. 5d)** and observed a significant increase in the promotion of PrLD SFPQ droplet formation with increasing concentrations of KIF5A in PEG3350 solution **(Fig. 5g)**. Analogous to the synergistic effect of KLC1 and *NORAD* in facilitating SFPQ PrLD LLPS, KIF5A could also cooperate with *NORAD* to promote the formation of SFPQ PrLD droplets **(Fig. 5h)**. We further purified Kinesin-1 **(Supplementary Fig. 5e)** composed of KIF5A and KLC1, and observed that Kinesin-1 alone formed aggregates **(Supplementary Fig. 5f)** and was insufficient to promote PrLD domain LLPS independently **(Supplementary Fig. 5g)**. Nonetheless kinesin-1 effectively promoted the formation of PrLD SFPQ droplets in the presence of *NORAD* **(Supplementary Fig. 5i, h)**. Collectively, our results demonstrated that *NORAD* not only independently promotes SFPQ LLPS but also serves as a scaffold that collaborates with KIF5A and KLC1 to enhance SFPQ LLPS **(Fig. 5j)**.

### Neuronal-specific *Norad* knockout mice exhibit axonal degenerative phenotype

To study the function of *Norad* in the nervous system, transgenic mice with neuronal-specific knockout of the 2900097C17Rik gene (the ortholog of *NORAD* gene in mouse) were generated (**Fig. 6A**). The genotyping of 2900097C17Rik KO mice and WT mice was verified by PCR **(Supplementary Fig. 6a, b)** where loxp was successfully inserted at both ends of exon I of the gene **(Supplementary Fig. 6f)**. We further performed spatiotesmporal transcriptomic analysis to revealed the expression patterns in WT and *Norad* KO groups **(Fig. 6d)**. As a result, *Norad* expression level was reduced in the KO group compared to the WT group in different areas of mouse brain **(Supplementary Fig. 6e)**. Furthermore, the expression levels of neuronal markers *Rbfox3* and *Actb* were notably reduced in KO mice as compared to those of WT litter mates **(Supplementary Fig. 6d),** indicating the total number of neurons was reduced by *Norad* KO. Moreover, in 8 out of 11 different brain regions, the proportion of neuronal cells was generally lower in KO groups compared to the WT groups, including the Cortical Subplate (CTXsp), Forebrain Structures (FBS), Hippocampal Formation (HPF), Hypothalamus (HY), Olfactory Area (OLF), Pallidum (PAL), Striatum (STR), Thalamus (TH), Ventral Lateral Nucleus (VL) **(Fig. 6e, f)**.

**Fig. 6:**
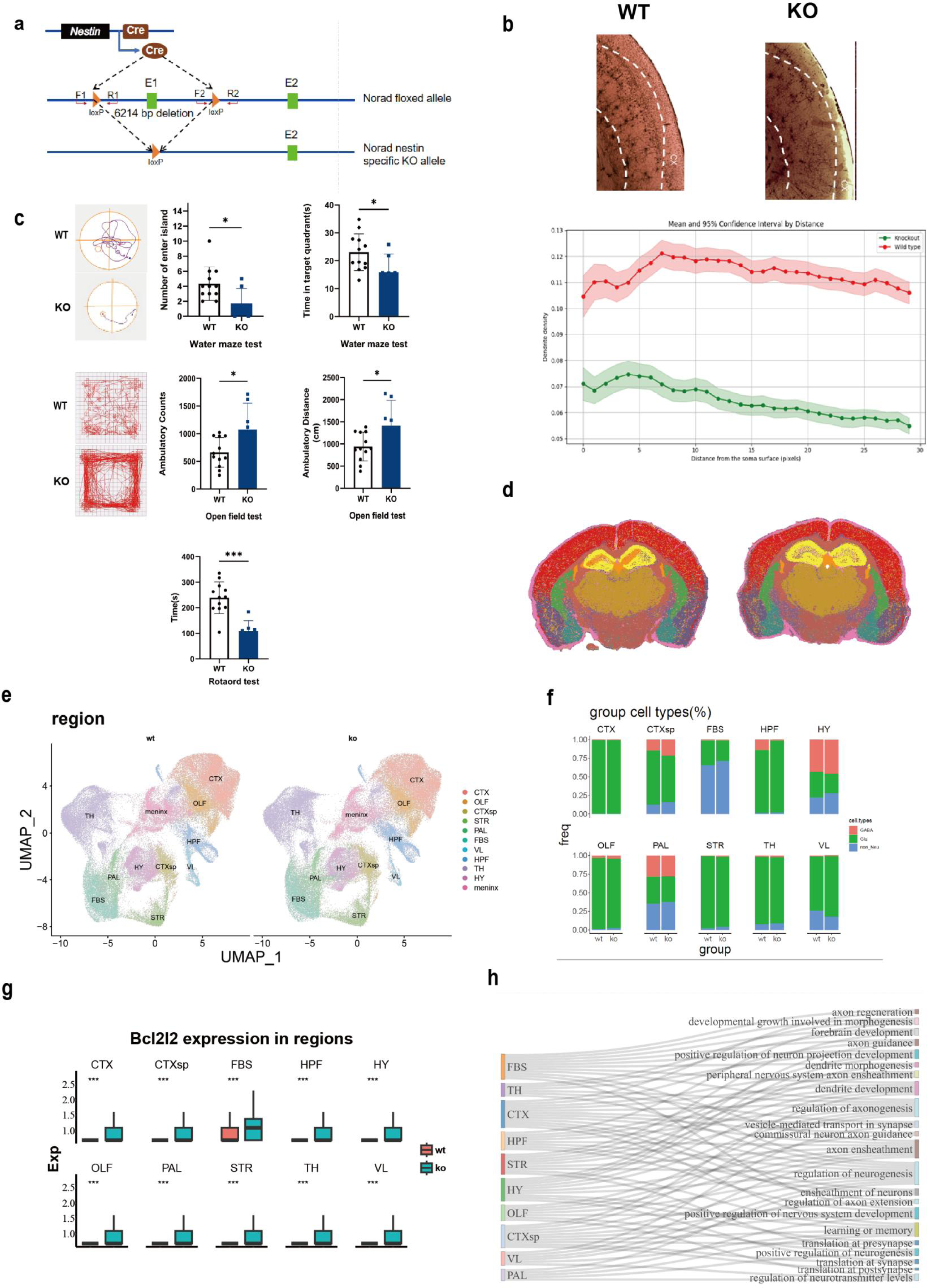
Neuronal-specific *Norad* knockout mice exhibit axonal degenerative phenotype. **a** Map of the *Norad* floxed allele. When *Norad^flox/flox^*mice are crossed with mice expressing Cre recombinase under an Nestin promotor, exons 1 of *Norad* are deleted. **b** Immunostaining on axonal marker Tuji-1, Golgi staining and the axon quantification in the brains of WT and KO mice. **c** Morris water maze test and open field experiments on WT and KO mice. **d** Spatiotesmporal transcritomic analysis in the brains of WT and *Norad* KO mice. **e** Distribution of cells in different brain regions of WT and *Norad* KO mice. **f** The proportion of neuronal cells including the Cortical Subplate (CTXsp), Forebrain Structures (FBS), Hippocampal Formation (HPF), Hypothalamus (HY), Olfactory Area (OLF), Pallidum (PAL), Striatum (STR), Thalamus (TH), and Ventral Lateral Nucleus (VL) in the brains of WT and *Norad* KO mice. **g** The expression of *Bcl2l2* mRNA in neuronal cells including the CTXsp, FBS, HPF, HY, OLF, PAL, STR, TH and VL of WT and *Norad* KO mice. **h** Sankey diagram for the 11 regions about differentially expressed genes in the brain of WT and *Norad* KO mice.

In neuronal axons, *Bcl2l2* mRNA is essential for axonal growth which is transported by KLC1/SFPQ^67,68^. However, spatial transcriptomics data revealed that in *Norad* KO group, the expression of *Bcl2l2* mRNA is accumulated in neural body compared with WT group, indicating depletion of *Norad* impairs the *Bcl2l2* mRNA transport **(Fig. 6g)**. These findings suggest that the defect of *Norad* in the brain results in disrupted transport of specific mRNAs and a decrease in the population of various neuronal cells, aligning with the phenotypes observed in SFPQ and KLC1 knockdown models ^69,70,71^. Moreover, the most common pathways shared by downregulated genes across the 11 regions are axon ensheathment and regulation of neurogenesis **(Fig. 6h)**. Based on these results, we propose an integrated transport regulatory model in which *Norad* mediates axonal transport , which is critical for proper neuronal viability.

To assess neuronal morphology, density, and distriution in the brain of WT and *Norad* KO mice, we performed immunostaining on axonal marker Tuji-1 and Golgi staining. The result indicates a significant reduction in the number of neurons in the hippocampus and cortex of 10-month-old mice after *Norad* knockout **(Fig. 6b and Supplementary Fig. 6c).** Furthermore, compared with WT mice, *Norad* KO mice showed increased movement distance and more standing times in open field experiments, significantly decreased spatial memory in water maze experiments, and significantly accelerated falling in rotarod test **(Fig. 6c)**. Collectively, neuronal-specific *Norad* KO mice showed significantly impaired motor function and decreased spatial memory ability. Together, these data provide compelling evidence that *Norad* brain-deficient animals results in neuronal axonal degenerative phenotype resembling axonal transport (my opinion is that there is little evidence to support axonal degeneration or axonal transport in thses mice, especially when compared with the findings in the hESC-neurons in the previous figures).

## Discussion

Our study elucidates a previously unrecognized mechanism by which the long non-coding RNA *NORAD* orchestrates axonal transport through liquid-liquid phase separation (LLPS), thereby maintaining neuronal integrity. We demonstrate that NORAD directly interacts with kinesin light chain 1 (KLC1) to promote the assembly of Splicing Factor Proline- and Glutamine-Rich (SFPQ) into dynamic LLPS condensates, enabling efficient cargo transport in axons. This discovery not only resolves the long-standing question of how specific RNAs regulate the SFPQ-KLC1 axis but also unveils a novel RNA-guided mechanism underlying axonal homeostasis.

The identification of *NORAD* as a direct interactor of KLC1 addresses a critical gap in understanding RNA-dependent axonal transport. Kinesins, a diverse group of microtubule-dependent motors, play a crucial role in facilitating the intracellular transport of various organelles^11,21,36^. While previous studies established that RNase-sensitive interactions mediate SFPQ-KLC1 binding^4,11^, the specific RNA species involved remained elusive. Our CRISPR-assisted RNA-protein interaction mapping (CARPID) and functional validation reveal *NORAD* as the key RNA orchestrating this process. This finding aligns with *NORAD*’s abundance in neuronal tissues^12,13,14^ and its evolutionary conservation, underscoring its biological significance. Furthermore, the decline of *NORAD* expression during aging and neurodegeneration^15,16^ suggests that its loss may contribute to axonal pathology, providing a mechanistic link between lncRNA dysfunction and age-related neurological disorders. Our CRISPR-assisted RNA-protein interaction mapping (CARPID) revealing 1075 common targets with >84% reproducibility across independent experiments, combined with functional validation showing complete rescue upon *NORAD* re-expression, strongly supports a direct mechanistic role rather than mere correlation.

The demonstration that *NORAD* promotes LLPS of SFPQ introduces a new paradigm for how RNA molecules facilitate cargo transport in crowded axonal environments. Axonal constraints,such as narrow diameters (often <2 μm)^26,27^ and high macromolecular density demand cargo flexibility to avoid physical crowding^24,25^. We show that SFPQ granules exhibit liquid-like properties, including fusion and rapid fluorescence recovery after photobleaching (FRAP), and that *NORAD* enhances these properties through its interaction with KLC1. This mechanism allows SFPQ condensates to deform and navigate axonal pathways efficiently, preventing stagnation and ensuring timely delivery of neuroprotective mRNAs like *Bcl2l2*. Notably, NORAD’s role in LLPS parallels its function in maintaining genomic stability via PUMILIO proteins^14^, suggesting a conserved mechanism across cellular contexts.

Our findings offer compelling insights into the pathogenesis of neurodegenerative diseases. *NORAD* depletion disrupts SFPQ granule dynamics, impairs mRNA localization, and induces axonal degeneration phenotypes reminiscent of aging-related neurodegeneration^16^. The motor deficits and reduced neuronal density observed in neuron-specific Norad knockout mice further underscore its physiological relevance. These results position *NORAD* and its associated LLPS machinery as potential therapeutic targets. For instance, enhancing *NORAD* expression or function could stabilize SFPQ condensates and restore axonal transport in conditions like Alzheimer’s disease or amyotrophic lateral sclerosis, where RNA dysregulation and transport deficits are prominent.

While our study provides mechanistic insights, several questions remain. First, the precise structural features of *NORAD* that enable KLC1 binding and LLPS promotion require further elucidation. Second, whether *NORAD* regulates other cargoes beyond SFPQ warrants investigation, given the diversity of RNAs and proteins identified in our CARPID analysis. Third, the temporal dynamics of *NORAD*-mediated LLPS in live neurons need higher-resolution imaging approaches to fully capture. Future studies could explore these aspects using advanced techniques such as super-resolution microscopy, structural biology, and high-throughput screening for *NORAD* mimetics or inhibitors.

A recent RNA-seq study of noncoding RNA expression in the subependymal zone of the human brain across different ages has reported a significant age-related decrease in NORAD expression^16^. These findings are particularly important given our new understanding of the role of NORAD in the neural system. Looking forward, our findings open several transformative research directions against age-related neurodegeneration. First , developing *NORAD*-based biomarkers for early detection of axonal degeneration. Second , engineering synthetic lncRNAs to modulate phase separation in neurodegenerative diseases. Third, exploring cross-talk between *NORAD* and other phase-separating systems in age-related pathologies. The convergence of lncRNA biology, phase separation physics, and neuroscience promises to revolutionize our understanding of neuronal maintenance in aging.

In summary, we reveal NORAD as a critical lncRNA that safeguards axonal integrity through LLPS-mediated transport. This work advances our understanding of RNA-guided biological processes and highlights the therapeutic potential of targeting NORAD in neurodegenerative diseases. As aging populations face increasing burdens of neurological disorders, unraveling such mechanisms becomes imperative for developing innovative treatments. We anticipate that this study will inspire future research into lncRNA-mediated phase separation and its role in neuronal health and disease.

## Methods

### Cell Culture

The hESC H1 was purchased from WiCell and maintained in DMEM/F12 (ThermoFisher) supplemented with essential eight (E8) (Thermo Fisher) in a Matrigel (Corning) coated (1:400 for 24h) polystyrene 2D culture system. Upon 80%–90% confluency, cells were dissociated in PBS with Accutase (Stemcell technologies) for 5-10 minutes at 37°C. Passaging was performed in 1:6 splitting ratios. For the first 24h after replacing, 10μM of ROCK inhibitor Y-27632 was included in the maintenance medium.

NSC induction and neuron differentiation were performed using the previously described canonical GSK3 and TGF-β receptor inhibitor inhibition protocol ^29^. For neural induction of hESCs, a previously published method was used with slight modifications, 4 μM CHIR99021, 3 μM SB431542, and 0.1 μM compound E (Stemcell technologies) were used in the induction medium from day 0 to day 7, after which the cells were cultured with 3 μM CHIR99021 and 2 μM SB431542 in neural induction medium. During passages, the ROCK inhibitor (Y-27632, 5 μM, Stemcell technologies) was used to promote cell survival. NSCs were expanded on Matrigel-coated surfaces. Spontaneous neuron differentiation was performed in DMEM/F12, 1xN2, 1xB27, 300 ng/mL cAMP (Sigma-Aldrich) and 0.2 mM vitamin C (Sigma-Aldrich) (referred to as differentiation media) on Matrigel and poly-L-ornithine (Sigma) coated surface. The single-cell spontaneous differentiation assay was carried out by plating 200 cells/well on 6-well plates in neural induction media supplemented with 3 μM CHIR99021 and 2 μM SB431542. After 3 d, the medium was switched to differentiation medium with 10 ng/mL BDNF and 10 ng/mL GDNF for another 14 d. For dopaminergic neuron differentiation, cells were first treated with 100 ng/mL SHH (C24II) and 100 ng/mL FGF8b in differentiation media for 10 d, and then with 10 ng/mL BDNF, 10 ng/mL GDNF, 10 ng/mL IGF1, 1 ng/mL TGF-β3 and 0.5 mM db- cAMP (Sigma-Aldrich) for another 14–21 d in differentiation media. For induction of motor neurons, cells were sequentially treated with 1 μM RA (Sigma-Aldrich) in differentiation media for 7 d, then with 100 ng/mL SHH (C24II) and 0.1 μM RA for an additional 7 d, and finally with 50 ng/mL SHH (C24II) and 0.1μM RA for another 7 d. The cells were terminally differentiated in the presence of 10 ng/mL BDNF and 10 ng/mL GDNF in the differentiation media for about 7 d. All growth factors were from R&D Systems. All tissue culture products were obtained from Invitrogen except where mentioned.

Cortical organoids performed using the previously described protocol ^30^. Feeder-free hESCs were seeded on Matrigel coated plates in TesR-E8 medium (Stemcell technologies) with daily medium changes for 4 days. After 90% confluence, the colonies were detached using Accutase (Stemcell technologies) in a 37° C incubator for 5 minutes and spun down at 200 x g for 3 minutes. The cell pellets were re- suspended in TesR-E8 medium with Dorsomorphin (1 uM) (Stemcell technologies) and SB431542 (10 uM) (Stemcell technologies). Then, the cells were transferred to the 6-well plate with ROCK inhibitor (5 uM) (Y-27632; Stemcell technologies) and kept in a shaker (75 rpm). After 3 days, the TesR-E8 medium was replaced with Neurobasal (Life Technologies) supplemented with GlutaMAX, B-27 (1%) (Gibco), N2 (1%) (Gibco), NEAA (1%) (Life Technologies), PS (1%) (LifeTechnologies), SB (10 uM) and Dorso (1 uM)] for 7 days. Subsequently, the spheres were cultured in Neurobasal with GlutaMAX, B-27 (1%), N-2(1%), and NEAA (1%), and PS (1%) added with FGF2 (20 ng/mL) (Life Technologies) for 7 days, followed by 7 extra days with the same medium supplemented with FGF2 (20 ng/mL) and EGF (20 ng/mL). For neural differentiation, the medium was additionally supplemented with BDNF (10 ng/mL), GDNF (10 ng/mL), NT-3 (10 ng/mL) (PeproTech), L-ascorbic acid (200 uM) and dibutyryl-cAMP (1 mM) (Sigma-Aldrich), which promoted maturation, and gliogenesis. The cerebral organoids were maintained in this medium with media changes every 3-4 days for as long as necessary.

### Generation of inducible-*NORAD* hESCs

The small guide RNAs (sgRNAs), which were targeted to the transcription start site (TSS) of *NORAD*, were inserted into the Pgl3.0 vector. The sgRNAs are shown in the Table. The Tre-Ef1α promoter-rTTA segment and BSD segment were cloned into the vector (BGI) and integrated into the TSS of *NORAD* with the designated sgRNAs respectively. 3μg of sgRNAs, 5μg of Cas9 plasmids, and 5μg of pLB donor vector were mixed and electroporated (Lonza) into hESCs. 1 day after electroporation, 2μg/ml Blasticidin S was added and individual colonies picked and expanded in 24- well plates. Genomic DNA was then purified using the TIANamp Genomic DNA Kit (DP304-03). The genomic integration was validated by PCR.

### Plasmid construction

The short hairpin RNAs (shRNAs) targeted to *NORAD* or KLC1 were inserted into the pLKO.1 vector (Addgene). The tet-on promoter fragments were cloned into the donor vector (introduced into the original promoter of *NORAD* locus of the hESCs).

PcDNA3.1-NORAD (Addgene) vector for *in vitro* transcription in RNA pull down assay. The cDNAs for the KLC1 ( NM_001130107) and SFPQ (NM_005066) genes used in this study were cloned into the PiggyBac and vector (Addgene). All constructed plasmids were verified by DNA sequencing. All primer sequences are listed in the Table .

### CARPID

Cells were washed twice using cold PBS and lysed with 500 μl lysis buffer (50 mM Tris-HCl (pH 7.4); 150 mM NaCl; 0.5% Triton X-100; 1 mM EDTA supplemented with fresh protease inhibitors (Roche)) at 4 °C for 10 min. The lysis was then spun down at 15,000 rpm. at 4 ° C for 10 min, separated 2% for input. After 2h of incubation at 4 ° C with rotation, biotinylated proteins were enriched with MyOne T1 streptavidin beads (Thermo Fisher) and washed 5 times with 500 μl ice-cold lysis buffer. Proteins were eluted from the beads by incubation into the elution buffer for 10 min at 95 ° C.

### Mass spectrometry analysis

The beads were suspended in 50 μl of elution buffer I (containing 50 mM Tris-HCl pH 8.0, 2 M urea, 10 µg ml–1 sequencing-grade trypsin from Thermo Fisher, and 1 mM DTT) and incubated at 30 °C with mixing at 400 r.p.m. for 60 min. Subsequently, the beads were eluted twice with 25 μl elution buffer II (comprising 50 mM Tris-HCl pH 8.0, 2 M urea, and 5 mM iodoacetamide). An additional 0.25 µg of trypsin was added to the combined eluates, followed by overnight incubation at 37 °C. The enzymatic digestion process was stopped by adding a 10% formic acid solution at a 1:25 (vol/vol) ratio. The digested samples were desalted using C18 tips from Thermo Fisher as per the manufacturer’s guidelines, and reconstituted in 20 μl of 0.1% formic acid. The analysis by liquid chromatography–tandem mass spectrometry (LC–MS/MS) was conducted using an Easy-nLC 1200 system connected to a Q Exactive HF mass spectrometer from Thermo Fisher.

### SpyCather protein expression and purification

For the recombinant purification, the SpyCatcher genes were transformed into *E. coli* Rosetta (DE3) pLysS competent cells. The single colony was selected and inoculated into a 25mL Luria-Bertani(LB) medium with 50 μg/ml kanamycin (Sigma Aldrich) at 37 °C. A final concentration of 1 mM isopropyl β-D-thiogalactoside (IPTG) (Sigma Aldrich) was introduced when the cells reached 0.8 at OD600. After incubating for 6 hours, cells were extracted and lysed in a lysis buffer (20mM Tris-HCl, pH 7.5, 500mM NaCl, 20mM imidazole) by sonification. The crude lysates were centrifuged at 18000 rpm for 30 min, then transferred into a HisTrap HP column (GE Healthcare) equilibrated with the binding buffer using the ÄKTA Prime instrument. After washing with 10 column volumes (CVs) of lysis buffer, the target protein was eluted with a gradient concentration of imidazole. The eluates were diluted to a final concentration of 50 mM NaCl and transferred into a column of anion ion exchange (GE Healthcare), and the ÄKTA used to collect the fractions. SDS-PAGE and Coomassie blue staining tested the purity of each fraction. Using 10-kDa molecular mass cut-off centrifugal filter unit, fractions that displayed relatively high purity were combined and concentrated. Purified proteins from SpyCatcher (8 mg / ml in PBS) were aliquoted and stored at-80 °C.

### SpyCatcher magnetic bead preparation

1 mg DynabeadsTM M-270 Epoxy(Invitrogen) were washed three times with 1 ml PBS then re-suspended in 50 μl PBS with 0.1 mg of SpyCatcher protein. Then washed three times with 1 ml Spybeads wash buffer (PBS containing 500 mM NaCl and 0.5 % Triton X-100) after a rotation at 25 ° C overnight followed by 6 hours blocking at 25 ° C in 1 ml of Spybeads blocking buffer (PBS containing 1 % BSA and 0.1 % Triton X-100). The SpyCatcher-coupled beads were stored at 4°C before use. Mass spectrometry was used to confirm the SpyCatcher protein covalently bound on beads after harsh washing sequentially with urea wash buffer (100 mM Tris, pH 7.4, 150 mM NaCl, 8 M urea, 0.2% Triton X-100), SDS wash buffer (100 mM Tris, pH 7.4, 150 mM NaCl, 1% SDS, 0.2% Triton X-100), high-salt wash buffer (50 mM Tris, pH 7.4, 2 M NaCl, 0.2% Triton X-100), and low-salt wash buffer (50 mM Tris (pH 7.4), 0.2% Triton X-100).

### SpyCLIP

The protocol has been described previously ^31^. Briefly, cells over-expressing SpyTag- SFPQ were irradiated at 400 mJ/cm^e^ in a UV Crosslinker (UVP, CL-1000) and lysed in lysis buffer (50 mM Tris (pH 7.4), 150 mM NaCl, 0.5% sodium deoxycholate, 0.1% SDS, 1% Triton X-100, and 1:100 protease inhibitor cocktail). Then the lysates were digested by RNase I (100 U/μl, Invitrogen) and immunoprecipitated by anti-FLAG magnetic beads. The 2’, 3’ cyclic monophosphate at the 3’ end of RNase I-treated RNAs was removed and ligased with Biotinylated 3’ adapter. Afterwards, RNP was released from FLAG beads using PSP (2 U/μl, GE Healthcare, 27-0843-01). The target proteins with associated RNAs were pull-downed by SpyCatcher. After Stringent washing, proteins were digested by Proteinase K (20 mg/ml, Roche) and RNAs released from SpyCatcher beads. Streptavidin beads were added to catch biotins at the 3’ end of the adaptor in the RNAs. On-bead reverse transcription was performed and cDNA generated. The cDNA was ligased with UMI 3’ adapter. Then, PCR amplification was performed using the same RP1 and RPI PCR primers as those used in the Illumina TruSeq Small RNA kit. The cDNA libraries were size-selected and purified in 6% polyacrylamide gel. The region between 160-180 bp was cut out and purified for sequencing, and performed by the Centre for Genomic Sciences, HKU.

The reads of the High-throughput sequencing of SpyCLIP were mapped to the human genome (version hg38) using the Bowtie 2 program for alignment. Mapped reads were normalized by bamCompare of the deepTools program. The generated bigwig files of the two clip samples and input were visualized on the Integrative Genomics Viewer (IGV).(fig.) The location (start site and end site) of the signals was defined by callpeaks function of MACS2. The binding motifs of SFPQ were identified by HOMER’s findMotifs program (-p 4 -rna -S 10 -len 6).

### Reverse transcription-quantitative PCR (RT-qPCR)

Total RNA was extracted from the cells using an RNAzol® RT (Molecular Research Center) according to the manufacturer’s instructions. Cells were homogenized with RNAzol and water. Then RNAs were precipitated using the same volume of isopropanol. The RNA pellets were washed with 70% ethanol and dissolved in water. 500 ng total RNA was used for synthesized cDNA with the PrimeScriptTM RT reagent Kit (Takara). The reverse transcribed cDNA was diluted 20-fold with water and 25 ng used in each qPCR reaction. The reactions were executed using SYBR Premix Ex Taq TM (Takara) on a LightCycler 480 (Roche) instrument. The GAPDH gene has been used as an internal Ctrl for gene expression analysis by qRT-PCR. The relative expression level was calculated using 2^-◻◻CT^. Each experiment was performed in triplicate.

### Western blotting

Cells were washed twice in ice-cold PBS and incubated in lysis buffer (150 mM NaCl, 50 mM Tris, pH 7.4, 1% Triton-X 100, 1 mM EDTA with a protease inhibitor cocktail (cOmplete EDTA-free, Roche)) for 15 min on ice. The lysates were pre-cleaned in a benchtop centrifuge for 10 min at full speed. Equal amounts of cell lysates were collected, mixed with 20 μl of 6 x sample buffer (0.35 M Tris-HCl pH 6.8, 10% SDS, 30% glycerol, 0.6 M DTT and 0.12% Bromphenol blue) and boiled at 95 °C for 10 min. Boiled samples were loaded into the electrophoresis of SDS-PAGE. After separation, the gels were transferred to a nitrocellulose (NC) filter membrane (Millipore). The membranes were blocked in 3% skimmed milk (Sigma) and incubated with primary antibody overnight at 4℃. The membranes were incubated with suitable secondary antibodies for 1 hour at room temperature after washing. The signals were visualized by enhanced chemiluminescence (ECL).

### Cytoplasmic and nuclear RNA fractionation

The cells were harvested, washed with cold PBS twice and re-suspended in 0.1% NP40. After homogenization, cell lysates were quick spun at 5000 g for 10 seconds. The supernatant was transferred into a new tube containing the cytoplasmic fraction. The pellet, which was a nuclear fraction, was gently washed twice with 0.1% NP40. The RNAs from cytoplasmic and nuclear fractions were extracted by RNAzol and subsequently analyzed by RT-qPCR.

### Co-immunoprecipitation

Neurons were harvested and washed with PBS. Re-suspended cells were lysed in lysis buffer. Ten percent of lysate was kept as input. The remaining supernatant was incubated with the corresponding antibody previously incubated with dynabeads Protein A or G at 4 °C overnight under shaking. The protein-bead complex was washed 5 times with a lysis buffer. For Western blotting, samples with SDS buffer were boiled at 95 ° C for 10 min.

### RNA immunoprecipitation

In each RIP assay, 3 μg antibodies were used for immunoprecipitation. Antibodies (Anti-KLC1 and SFPQ, ab174273; ab38148 ) were used in this study for RNA enrichment associated with the immunoprecipitated proteins. The RNAzol kit was used to extract the co-immunoprecipitated RNAs, using the PrimeScriptTM RT reagent Kit (Takara), cDNA was synthesized and then analyzed by qPCR.

### RNA Pulldown

*NORAD* fragments were amplified with primers containing T7 promoter sequences (Table 2.4). MEGAscript T7 Transcription Kit (Ambion) was used for the Biotin RNA labeling. *In vitro* transcribed RNA was treated with DNase I and purified with an RNeasy kit (Qiagen). 30 pmol purified biotinylated RNA was heated to 90°C in 60 µL RNA structure buffer (10 µM Tris-Cl pH 7.0, 0.1 M KCl, 10 mM MgCl2) for 2 minutes then put on ice for 2 minutes. 2×10^7^ cells were harvested by scraping and snap-frozen prior to resuspension in 1.2 mL lysis buffer [150 mM NaCl, 50 mM Tris-Cl pH 7.5, 0.5% Triton X-100, 1mM PMSF, 1x protease inhibitor cocktail (Roche), and 100 U/ml of SUPERaseIN (Ambion)]. Lysates were sonicated using a Bioruptor (Diagenode) for 10 min with 30 sec on/off cycles and pre-cleared with 50 µL washed streptavidin C1 Dynabeads (Invitrogen) at 4 °C for 1 hour. 30 pmol biotinylated RNA was then added to pre-cleared lysates and rotated at 4 °C for 2 hours. Beads were washed 5 times with lysis buffer at 4°C and re-suspended in SDS sample buffer. After boiling at 95 °C for 10 min, the samples were subsequently analyzed by Western blot.

### RNA FISH and Immunofluorescence assay (IF)

Cells were seeded on 8-well Millcell EZ slides (Millipore) coated with Poly-lysine (Sigma) and allowed to attach and grow for varying periods of time. The slides were fixed with 4% paraformaldehyde (PFA) for 15 min and washed with PBS three times, then fixed cells were permeabilized with 0.25% Triton-X 100 for 10 min, and washed with PBS three times. Then Stellaris® RNA probes (Table 2.6) for *NORAD* and various primary antibodies were diluted in hybridization buffer at a 1:100 ratio diluted in FISH washing buffer (10% formamide and 10% SSC in H_2_O) added into cells. The slides were incubated at 37 °C for 4-5 h in the dark. After washing with FISH washing buffer, cells were incubated with corresponding secondary antibodies at room temperature for 30 min in the dark. After washing, the slides were mounted with anti- fade fluorescent mounting medium containing DAPI (VECTASHIELD). The images were captured using an LSM 780 confocal microscope, and proprietary ZEN software used for data analysis.

### Protein expression and purification

*Escherichia coli* BL21-CodenPlus (DE3)-RIPL chemically competent cells (WeidiBio) were transformed with His10-MBP-TEV-PrLD-EGFP, His10-MBP-TEV-RRM- EGFP, His10-MBP-TEV-CTD-EGFP, and His10-MBP-TEV-△PrLD-EGFP plasmids, respectively. Bacteria were grown to an optical density of 0.8-1.0 and proteins were induced with 1 mM IPTG, after which bacteria were grown for an additional 18 h at 16 °C. Bacteria were collected by centrifugation and washed with lysis buffer containing 100 mM HEPES (pH 7.5), 2 M NaCl, 20 mM imidazole, 2 mM TCEP, 0.5 mM PMSF, 5% glycerol, and EDTA-free Protease Inhibitor Cocktail (MedChemExpress) to remove residual LB medium. Washed bacteria were further resuspended in lysis buffer and lysed by sonication. Cell pellets were cleared by centrifugation (20,000g × 30 min, 4 °C). The supernatant was then incubated with Ni-NTA beads (GE Health) and washed with 10 column volumes of lysis buffer (without PMSF, and Protease Inhibitor Cocktail). After an additional wash with 10 column volumes of Wash Buffer 1 containing 50 mM HEPES (pH 7.5), 2 M NaCl, 40 mM imidazole, 2 mM TCEP, and 5% glycerol, 5 column volumes of Wash Buffer 2 containing 50 mM HEPES (pH 7.5), 2 M NaCl, 40 mM imidazole, 2 mM TCEP, and 5 column volumes of Wash Buffer 3 containing 50 mM HEPES (pH 7.5), 150 mM NaCl, 40 mM imidazole, 2 mM TCEP, protein was eluted with elution buffer containing 50 mM HEPES (pH 7.5), 150 mM NaCl, 200 mM imidazole, 2 mM TCEP. Protein eluent fractions were pooled and concentrated via centrifugal filtration (Amicon Ultra Centrifugal Filter Unit, 30K MWCO) to 100 μl. After an additional centrifugation step (15,000g × 10 min, 4 °C), protein solution was further purified with SEC buffer containing 50 mM HEPES, pH7.5, 150 mM NaCl, 1 mM TCEP by size exclusion chromatography via Superdex 200 Increase Column (24 mL) mounted on AKTA Pure (GE Health). Proteins were further concentrated via centrifugal filtration (Amicon Ultra Centrifugal Filter Unit, 30K MWCO), then flash-frozen and stored at −80 °C.

Human HEK293F cells were transfected with MBP-TEV-mCherry-Kif5A and 3 × Flag-KLC1 plasmids simultaneously at a cell count range 2 × 10^6^/mL to 3 × 10^6^/mL. Cells were collected and washed after 48 h incubation by centrifugation. Washed cells were further resuspended and lysed by incubation in lysis buffer containing 1 × PBS, 1% Triton X-100, 5 mM ATP, 5 mM MgCl_2_, 2 mM TCEP, 0.5 mM PMSF, and EDTA-free Protease Inhibitor Cocktail (MedChemExpress) upon moderate shaking for 30 min at 4 °C. Cell pellets were cleared by centrifugation (20,000g × 30 min, 4 °C). The supernatant was then incubated with Anti-DYKDDDDK G1 Affinity Resin for 4 h (GeneScript). Then the affinity resin was washed with 50 column volumes of Wash Buffer 4 containing 1 × PBS, 2 mM TCEP, 5 mM ATP, 5 mM MgCl_2_ and 50 column volumes of Wash Buffer 5 containing 1 × PBS, 1 mM TCEP. Proteins were eluted by 200 mM 3 × DYKDDDDK peptide dissolved in Wash Buffer 5, concentrated via centrifugal filtration (Amicon Ultra Centrifugal Filter Unit, 30K MWCO), flash-frozen and stored at −80 °C.

### *In vitro* phase separation assays

Purified proteins were diluted in phase separation buffer containing 50 mM HEPES, pH7.5, 150 mM NaCl, 1 mM TCEP, 1 U TEV protease, and 10% PEG3350 (optional). For RNA-induced phase separation experiments, *in vitro*-transcribed RNAs labeled with Cyanine 3-UTP were diluted in water and denatured at 70 °C for 10 min before being added to protein mixtures in phase separation buffer. Protein or protein-RNA solutions were incubated at room temperature in an RNase free 384-well glass bottom plate (Cellvis) for 30 - 60 min before imaging.

### Axon degeneration assay *in vitro*

Compartmented chamber cultures were fixed at room temperature with 4% PFA diluted 1:2 in PBS for 10 min. DRGs were permeabilized with 0.1% Triton X-100 for 10 min, blocked with 3% BSA in 0.1% Triton X-100 for 1 h at room temperature, and incubated with rabbit anti-Tuj1 overnight at 4 °C. Cultures were then incubated with secondary antibodies (1:1,000; Invitrogen) for 1 h at room temperature. Images of distal axon tips were obtained using a×40 oil objective, and axonal degeneration quantified using the method described by Sasaki et al. (2009)^66^. TUJ1-stained images were binarized using ImageJ to convert axonal areas to black and background areas to white. To detect fragmented (degenerating) axons, the particle analyzer function of Image J software was used (for total area we analyzed particle size, 0-infinity; and for axon fragments, we set the particle size, 0-1,000). The degeneration index was calculated as the ratio of the area of fragmented axons to the total axon area.

### Dendrite density in mice brain slides

To compare dendrite distributions in same brain regions between knockout (KO) and wild-type (WT) groups, we developed a Napari-based annotation tool for semi-automated soma center identification and dendrite extraction. Users begin by loading an 8-bit TIFF image and adjusting processing parameters, including intensity threshold, size constraints, and Frangi filter settings. The software then detects soma centers based on morphological properties and extracts dendrites using a skeletonized Frangi filter response. Since it is not possible to precisely label all dendrite fibers, we adopt a selective annotation strategy: users manually choose somas where dendrites are well extracted by interactively adding or removing soma points through clicks.

This ensures that only high-confidence regions contribute to the analysis. The final annotations, including soma centers and dendrite skeletons, are saved for further analysis. We computed the mean and 95% confidence interval (CI) of dendrite density across distances from the soma surface. For each distance bin, we applied an independent two-sample t-test to assess significant differences between groups. The results were visualized as mean density curves with shaded confidence intervals.

### Statistics and reproducibility

Unless otherwise indicated, Student’s t test or two-way analysis of variance (ANOVA) with the Bonferroni post-test were used in this study. P values are represented as *: * p<0.05, ** p<0.01,*** p<0.001, **** p<0.0001. Graphpad Prism 6 was used to plot graphs and to perform statistical analysis.

### Data availability

All data needed to evaluate the conclusions of this Article are presented in supplementary materials. The source of CARPID data has been uploaded to the public database Figshare and can be accessed using the following Digital Object Identifier (DOI): 10.6084/m9.figshare.28661018.

## Acknowledgements

The research was in part supported by the RGC GRF (17124921), the Collaborative Research Fund (C1024-22G), the ITF fund (ITS/087/22) and National Key Research and Development Program of China (2023YFA0914904); Program Shenzhen Science and Technology Innovation Programme (SGDX2024011511310000) . J.H. thanks the L & T Charitable Foundation, the Program for Guangdong Introducing Innovative and Entrepreneurial Teams (2019BT02Y198) and Shenzhen Key Laboratory for Cancer Metastasis and Personalized Therapy (ZDSYS20210623091811035) for their support.

## Author Contributions

S.H., H.C., J.D.H.and Q. L designed the experiments and interpreted the results. S.H., H.C., J.L., Y.W., S.C.,P.S , Z.X ,R.Z. F.D, L. R. W.Y . K.A.B ,J.Y ,R.L,,J.X ,J.Q .G M.,,K.L ,C.L ,and P.Z performed the experiments and data analysis. J.J , Z.Q , C. L ,P.Z edited the manuscript and provided comments. S.H., H.C., Y.W., J.D.H. and Q.L wrote and revised the manuscript. J.D H. and Q. L co-supervised the whole project and acquired financial support for the project.

## Additional information

Correspondence and requests for materials should be addressed to Jian-Dong Huang and Qizhou Lian.

---

**Supplementary Figure 1.**
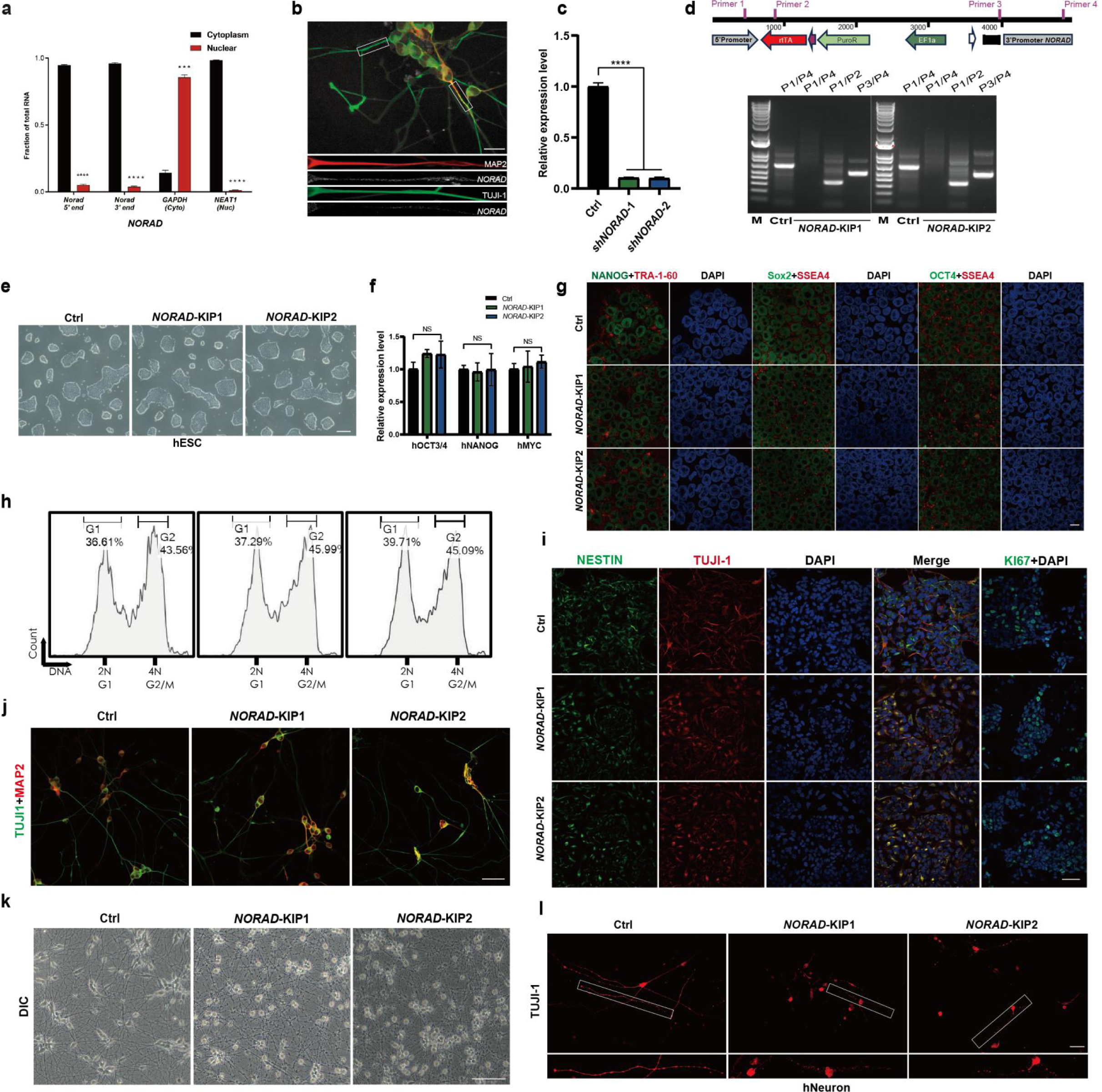
Depletion of *NORAD* results in axonal degeneration. **a** Subcellular fractionation in hESC-derived neuron followed by qPCR using primers located at the 3′ or 5′ end of NORAD, GAPDH was cytoplasmic Ctrl and NEAT1 was nuclear Ctrl. n=3 biological replicates each with three technical replicates. (mean ± SEM, n = 3, biological replicates; \*\*\**p* < 0.001, \*\*\*\**p* < 0.0001, one-way ANOVA). **b** Representative *NORAD* RNA FISH (white) images of hESC-derived neuron. TUJI- 1 (green), MAP2 (red). Scale bar: 20 μm. **c** qRT-PCR analysis of two distinct shRNAs targeting *NORAD* (sh*NORAD-1* and sh*NORAD-2*) in neurons. (mean ± SEM, n = 3, biological replicates, \*\*\*\**p* < 0.0001, one-way ANOVA). **d** PCR validation of Tet-on promotor knock-in in *NORAD*-KIP1/2 hESCs. Upper: scheme of amplified genomic sequence using primer sets. Lower: genotyping by agarose gel electrophoresis (2% agarose). **e** The colony morphology of the hESCs in Ctrl and knockdown groups. Scale bar: 200 μm. **f** The mRNA expression of OCT4, NANOG and MYC in the *NORAD*-KIP1/2 hESCs. (mean ± SD, n = 3, biological replicates; n.s., not significant, one-way ANOVA). **g** Pluripotency of Ctrl or *NORAD*-KIP1/2 hESCs. Immunostaining of NANOG (green)/ TRA-1-60 (red)(lane 1), SOX2 (green)/ SSEA4 (red) (lane3), OCT4 (green)/ SSEA4 (red) (lane 5), and nucleus (DAPI, blue). Scale bar: 20 μm. **h** Cell cycle analysis of Ctrl and *NORAD*-KIP1/2 hESCs. **i** Immunostaining of Ctrl or *NORAD*-KIP1/2 NSCs using antibodies against Nestin (green), TUJI-1 (red), and KI67 (green). Scale bar: 50 μm. **j** TUJI-1 immunostaining (green) and Map2 (red) in Ctrl and *NORAD*-KIP1/2 neurons. Scale bar: 50 μm. **k** The morphology of Ctrl and *NORAD*-KIP1/2 neurons. Scale bar: 100 μm. **l** TUJI-1 immunostaining (red) in Ctrl and *NORAD-*KIP1/2 neurons. Scale bar: 50 μm.

**Supplementary Figure 2.**
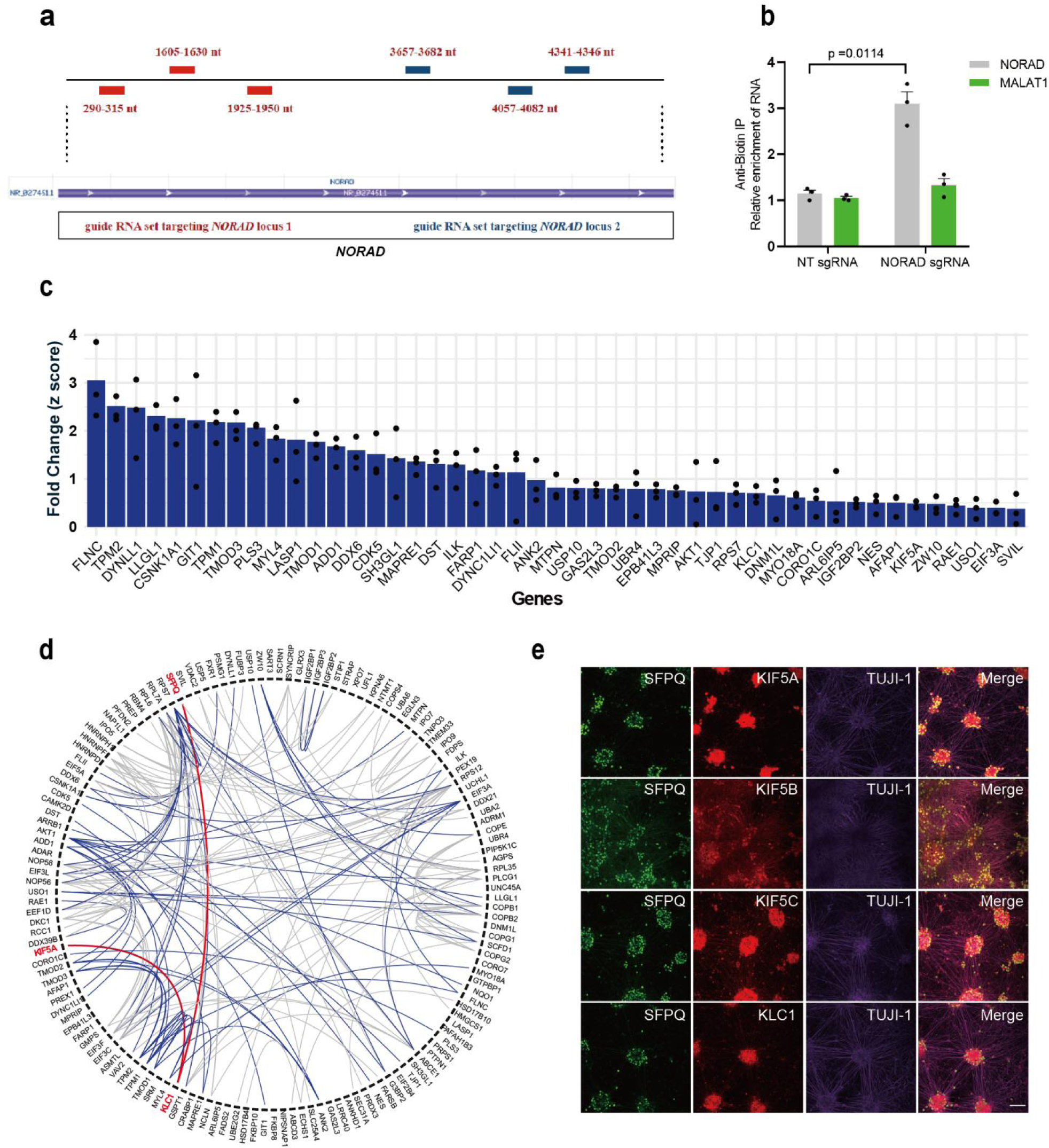
Identification of proteins interacting with *NORAD* by CARPID. **a** The location of the sets of gRNAs on *NORAD*. **b** The specificity of gRNA targeting *NORAD* was confirmed using RIP-qPCR with a biotin antibody, with MLAT1 utilized as a Ctrl. (mean ± SEM, n = 3, biological replicates; p = 0.0114, one-way ANOVA). **c** Log2 LFQ of cytoskeleton-localized *NORAD*-interacting proteins. The column represents the mean of three biological replicate experiments (individual data points). **d** Predicted interactions among enriched *NORAD*-interacting proteins. The chord diagram demonstrates the STRING analysis of upregulated *NORAD*-interacting proteins with high confidence (STRING confidence > 0.7). Chords represent a predicted interaction between two proteins. Blue chords indicate the involvement of cytoskeleton-localized proteins. Red chords indicate KLC1-interacting proteins. **e** Staining of endogenous SFPQ, KIF5s and TUJI-1 in hESc-derived neurons. SFPQ (green), KIF5 (red), TUJI-1 (purple). Scale bar: 100 μm.

**Supplementary Figure 3.**
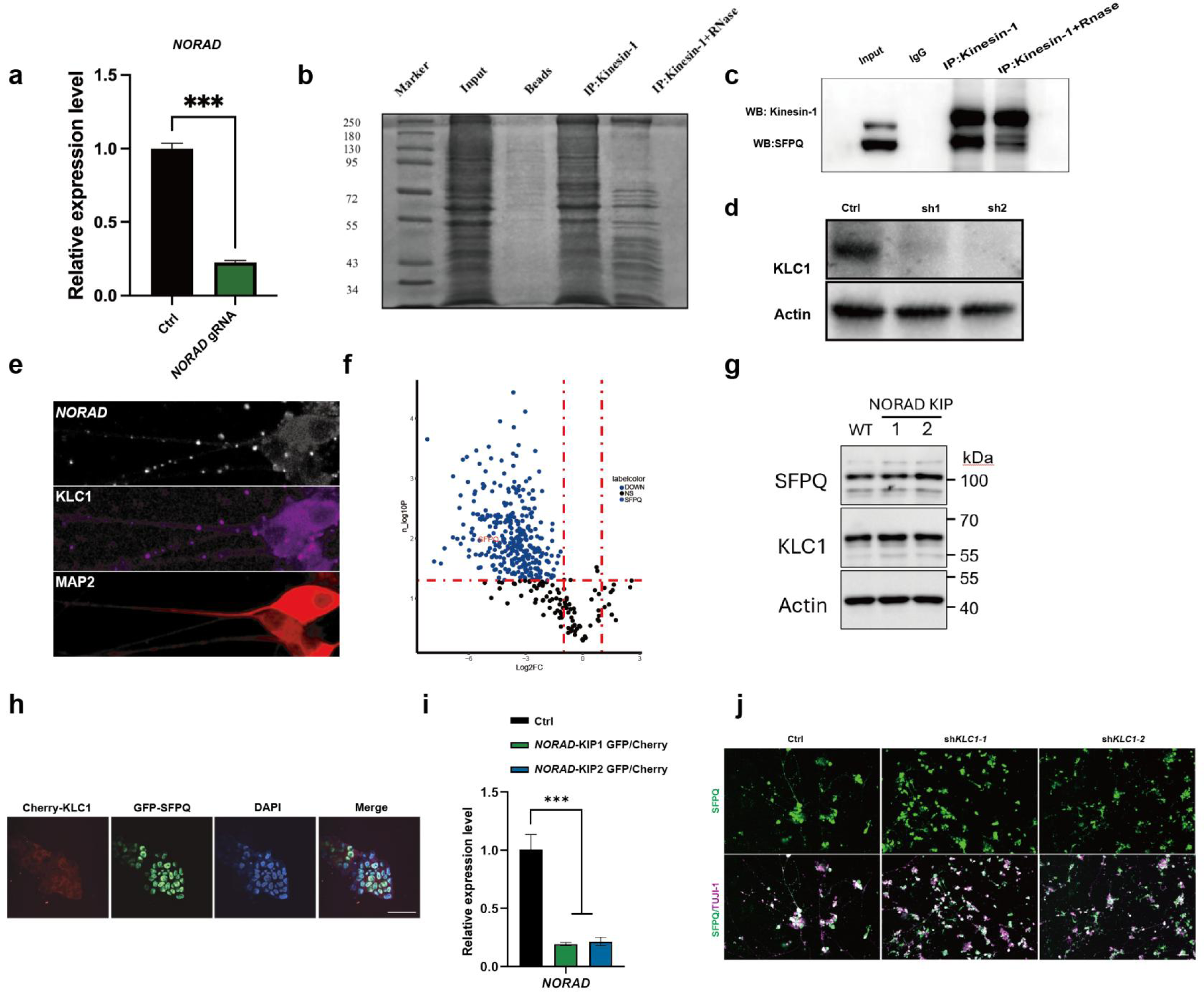
*NORAD* regulates the intracellular transport of SFPQ by binding with KLC1.**a** Expression of *NORAD* was analyzed by RT-qPCR in Ctrl and CRISPR-Cas13 knockdown neurons. (mean ± SEM, n = 3, biological replicates, \*\*\**p* < 0.001, one- way ANOVA). **b** SDS-PAGE analysis followed by Coomassie staining of KIF5B Co- IP with or without RNase treatment. Anti-rabbit normal IgG was used as the negative Ctrl. **c** HEK 293T lysates were treated with or without RNase followed by Kinesin-1 Co-IP. **d** Expression of KLC1 analyzed by western blot in hESC. **e** Representative *NORAD* RNA FISH (white) images of hESC-derived neuron. KLC1 (purple), MAP2 (red). Scale bar: 10 μm. **f** Volcano plot of MS of KLC1 immunoprecipitation of neuronal cells lysates treated with or without RNase. **g** Western blots of SFPQ or KLC1 in WT and *NORAD* KIP1/2 neuron. **h** Fluorescence of GFP-SFPQ (green) and mCherry-KLC1 (red) in hESC. Scale bar: 50 μm. **i** qRT-PCR analysis of *NORAD* expression in *NORAD*-KIP1/2 Cherry/KLC1 hESCs. (mean ± SEM, n = 3, biological replicates, \*\*\**p* < 0.001, one-way ANOVA). **j** Immunostaining SFPQ (green) and TUJI-1 (purple) in Ctrl and sh*KLC1* neurons. Scale bar: 100 μm.

**Supplementary Figure 4.**
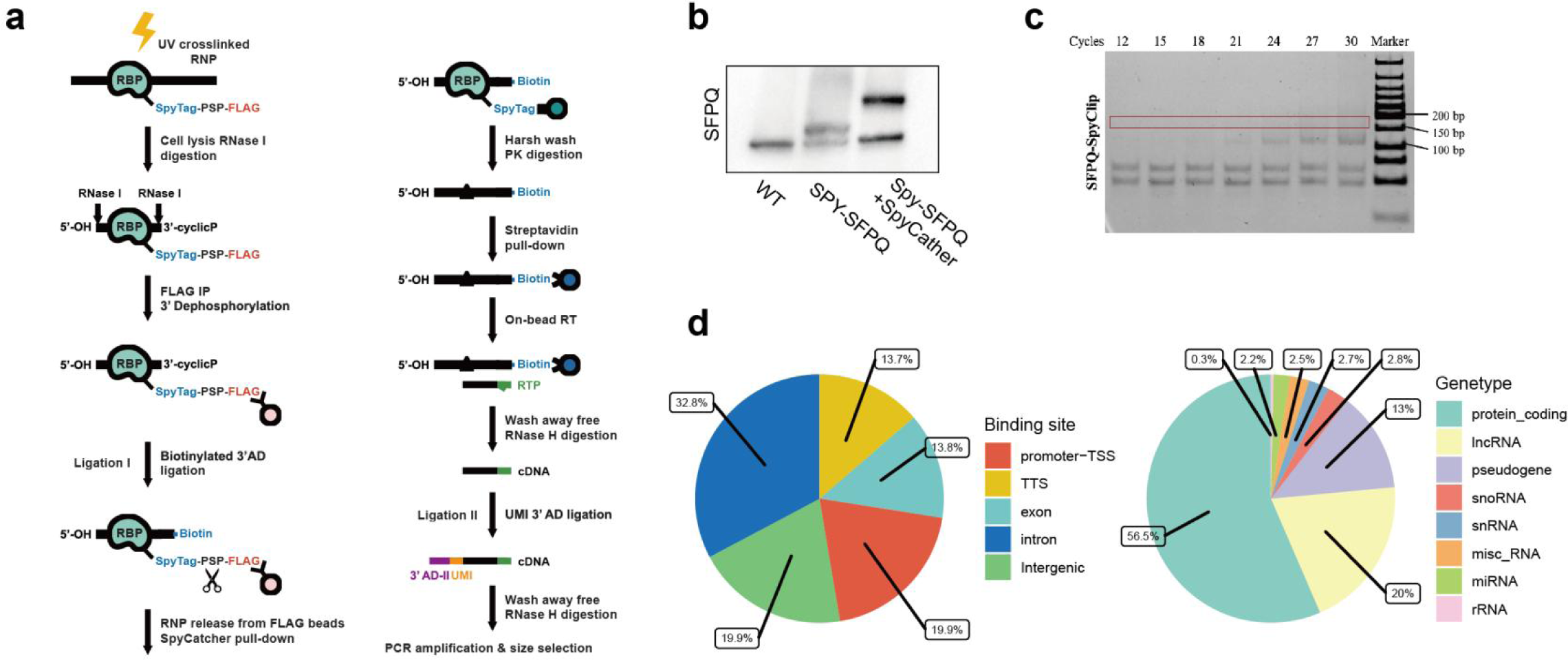
*NORAD* facilitates axonal transport of SFPQ via liquid liquid phase separation. **a** Diagrammatic illustration of the SpyCLIP experiment. **b** The modified SpyCatcher protein efficiently formed a stable complex with Spy-tagged SFPQ. **c** Cycles present a reasonable amount of cDNA for deep sequencing, and 27 PCR cycles were selected for library preparations. For deep sequencing, the region identified by the rectangle (160 - 180 bp) was recovered. **d** The distribution of the SFPQ Spy-CLIP peaks in the human genome. TSS transcription start site, TTS transcription termination site (left) and other binding sites identified in various RNA-species (Right).

**Supplementary Figure 5.**
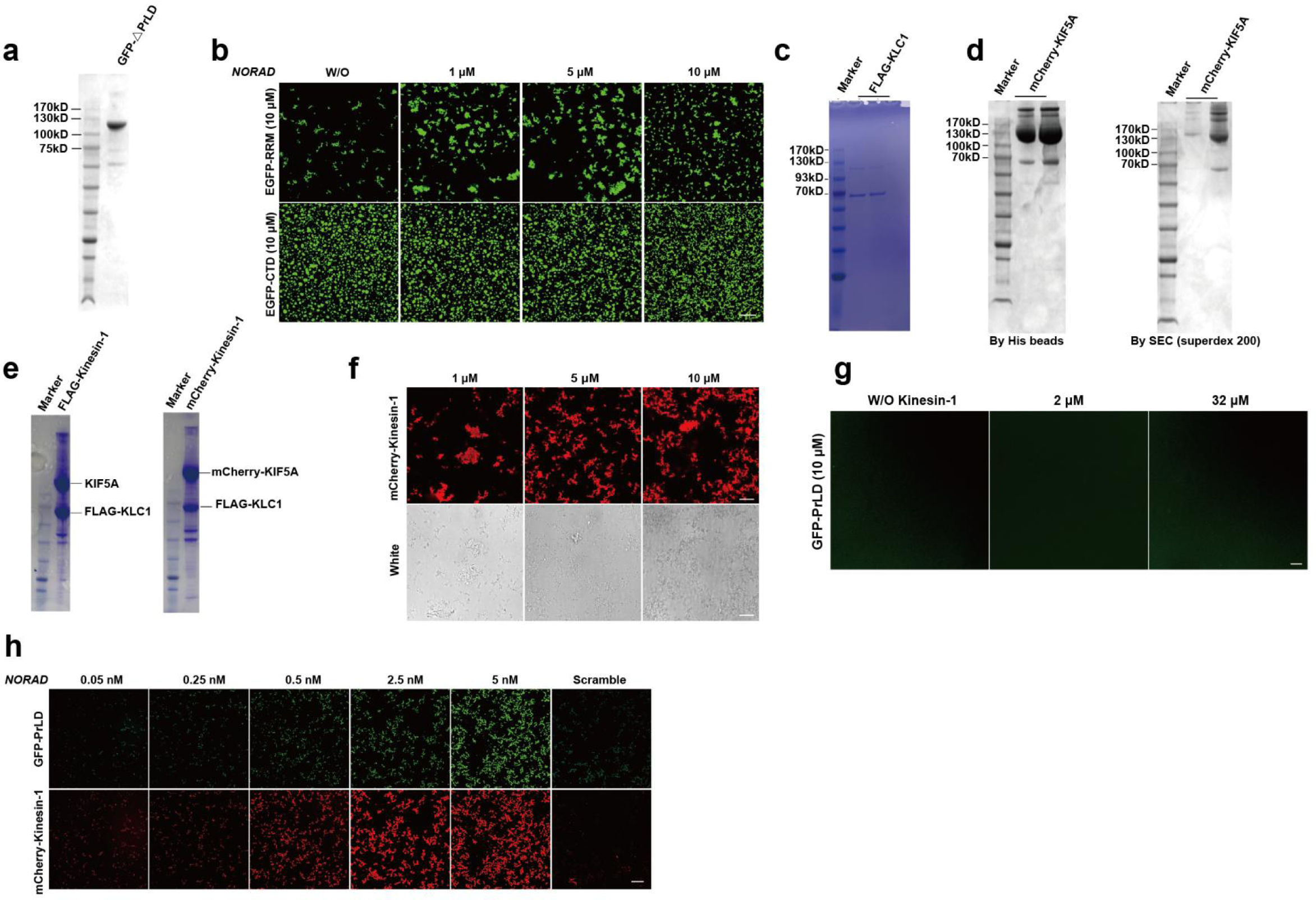
*NORAD* collaborates with Kinesin-1 to promote the LLPS of SFPQ-PrLD domain. **a** Proteins purifications of GFP-ΔPrLD domain. **b** Confocal images of EGFP-CTD domain and RRM domian droplets with different concentrations of *NORAD*. Scale bars: 20 μm. **c-e** Proteins purifications of Flag-KLC1 (C), mCherry-KIF5A (D) and Kinesin-1 protein with FLAG and mCherry tag expression trials in *Escherichia coli* BL21 (DE3). **f** Confocal images of mCherry-Kinesin-1 (red) at varying concentrations (top panels) alongside corresponding DIC images (bottom panels). Scale bars: 20 μm. **g** Confocal images of GFP-PrLD of SFPQ (10 μM) with or without Kinesin-1 (2 μM or 32 μM). Scale bars: 10 μm. **h** Confocal images of 10 μM GFP-PrLD (green) droplet with different concentration of mCherry-KIF5A (red) and *NORAD*. Scale bars: 20 μm.

**Supplementary Figure 6.**
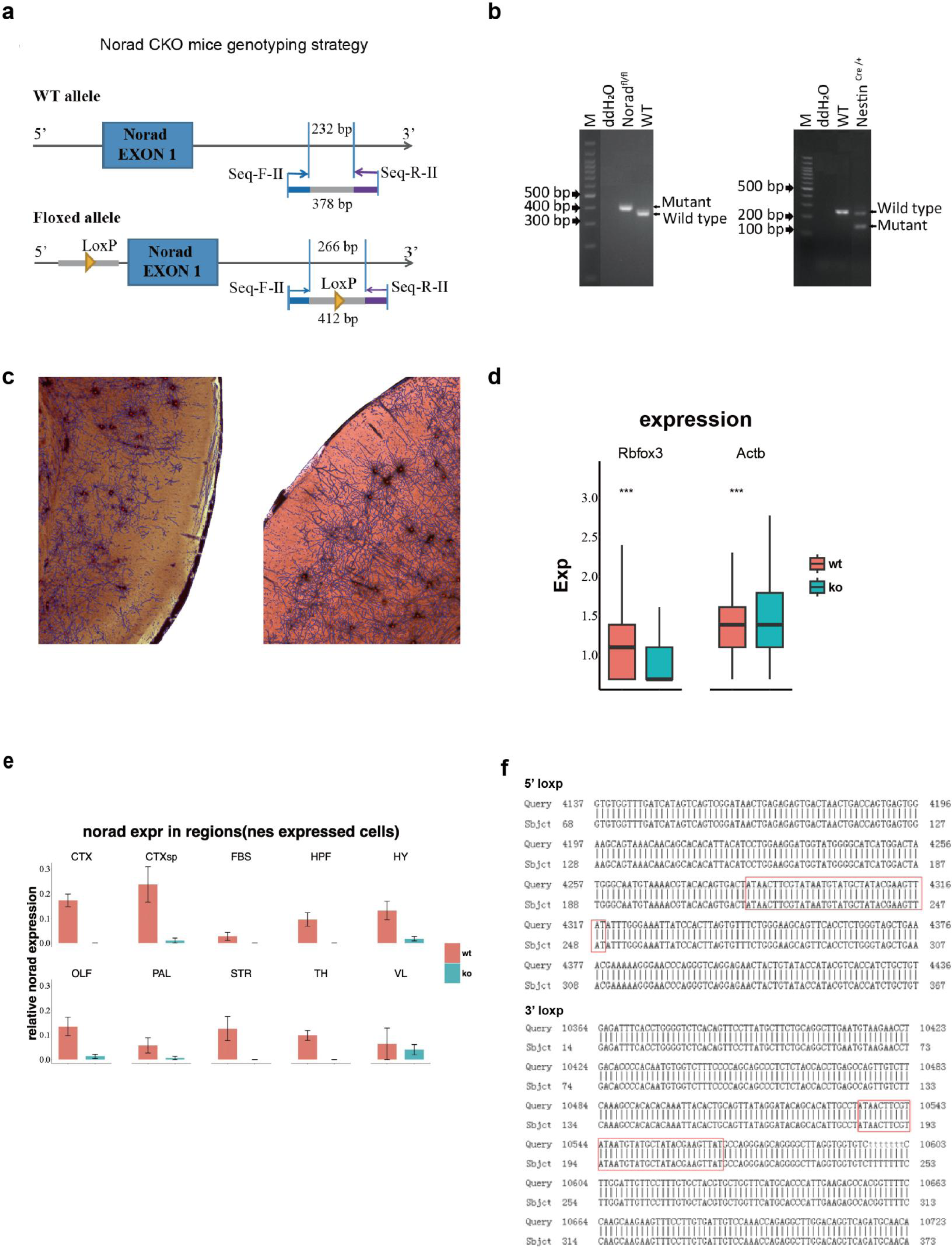
Neuronal-specific Norad knockout mice exhibit axonal degenerative phenotype. **a** Map of the *Norad* floxed allele and primer. **b** PCR genotyping of the *Norad* allele. Representative agarose gel showing amplicons from WT and Mutant mice. **c** Immunostaining on axonal marker Tuji-1 and Golgi staining in the brains of WT and KO mice. **d** The *Rbfox3* and *Actb* expression in KO mice as compared to those of WT litter mates. **e** The relative *Norad* expression in the 10 regions of WT and *Norad* KO mice. **f** Alignment of the *Norad* genomic region confirming correct insertion of the loxP sites.

## References

1. Holt CE, Martin KC, Schuman EM. Local translation in neurons: visualization and function. Nature structural & molecular biology. 2019;26(7):557–566.

2. Sahoo PK, Smith DS, Perrone-Bizzozero N, Twiss JL. Axonal mRNA transport and translation at a glance. Journal of cell science. 2018;131(8):jcs196808.

3. Cosker KE, Fenstermacher SJ, Pazyra-Murphy MF, Elliott HL, Segal RA. The RNA-binding protein SFPQ orchestrates an RNA regulon to promote axon viability. Nat Neurosci. May 2016;19(5):690–696. doi:10.1038/nn.4280

4. Fukuda Y, Pazyra-Murphy MF, Silagi ES, et al. Binding and transport of SFPQ-RNA granules by KIF5A/KLC1 motors promotes axon survival. Journal of Cell Biology. 2021;220(1)

5. Taylor R, Hamid F, Fielding T, et al. Prematurely terminated intron-retaining mRNAs invade axons in SFPQ null-driven neurodegeneration and are a hallmark of ALS. Nature Communications. 2022;13(1):6994.

6. Cioni J-M, Lin JQ, Holtermann AV, et al. Late endosomes act as mRNA translation platforms and sustain mitochondria in axons. Cell. 2019;176(1):56–72. e15.

7. Cosker KE, Pazyra-Murphy MF, Fenstermacher SJ, Segal RA. Target-derived neurotrophins coordinate transcription and transport of bclw to prevent axonal degeneration. Journal of Neuroscience. 2013;33(12):5195–5207.

8. Courchesne SL, Karch C, Pazyra-Murphy MF, Segal RA. Sensory neuropathy attributable to loss of Bcl-w. Journal of Neuroscience. 2011;31(5):1624–1634.

9. Pease-Raissi SE, Pazyra-Murphy MF, Li Y, et al. Paclitaxel reduces axonal Bclw to initiate IP3R1-dependent axon degeneration. Neuron. 2017;96(2):373–386. e6.

10. Huang J-D, Brady ST, Richards BW, et al. Direct interaction of microtubule-and actin-based transport motors. Nature. 1999;397(6716):267–270.

11. Saez TM, Fernandez Bessone I, Rodriguez MS, et al. Kinesin-1-mediated axonal transport of CB1 receptors is required for cannabinoid-dependent axonal growth and guidance. Development. 2020;147(8):dev184069.

12. Lee S, Kopp F, Chang T-C, et al. Noncoding RNA NORAD regulates genomic stability by sequestering PUMILIO proteins. Cell. 2016;164(1-2):69–80.

13. Elguindy MM, Kopp F, Goodarzi M, et al. PUMILIO, but not RBMX, binding is required for regulation of genomic stability by noncoding RNA NORAD. Elife. Jul 25 2019;8doi:10.7554/eLife.48625

14. Elguindy MM, Mendell JT. NORAD-induced Pumilio phase separation is required for genome stability. Nature. 2021;595(7866):303–308.

15. Kopp F, Elguindy MM, Yalvac ME, et al. PUMILIO hyperactivity drives premature aging of Norad-deficient mice. Elife. 2019;8:e42650.

16. Barry G, Guennewig B, Fung S, Kaczorowski D, Weickert CS. Long non-coding RNA expression during aging in the human subependymal zone. Frontiers in neurology. 2015;6:45.

17. Wang C, Duan Y, Duan G, et al. Stress induces dynamic, cytotoxicity-antagonizing TDP-43 nuclear bodies via paraspeckle LncRNA NEAT1-mediated liquid-liquid phase separation. Molecular cell. 2020;79(3):443–458. e7.

18. Saito M, Hess D, Eglinger J, et al. Acetylation of intrinsically disordered regions regulates phase separation. Nature chemical biology. 2019;15(1):51–61.

19. Lin Y, Protter DS, Rosen MK, Parker R. Formation and maturation of phase-separated liquid droplets by RNA-binding proteins. Molecular cell. 2015;60(2):208–219.

20. Lee M, Sadowska A, Bekere I, et al. The structure of human SFPQ reveals a coiled-coil mediated polymer essential for functional aggregation in gene regulation. Nucleic Acids Res. Apr 20 2015;43(7):3826–40. doi:10.1093/nar/gkv156

21. Kanai Y, Dohmae N, Hirokawa N. Kinesin transports RNA: isolation and characterization of an RNA-transporting granule. Neuron. 2004;43(4):513–525.

22. Thivierge C, Bellefeuille M, Diwan S-S, et al. Paraspeckle-independent co-transcriptional regulation of nuclear microRNA biogenesis by SFPQ. Cell Reports. 2024;43(9)

23. Ripin N, Parker R. Formation, function, and pathology of RNP granules. Cell. 2023;186(22):4737–4756.

24. Sabharwal V, Koushika SP. Crowd control: effects of physical crowding on cargo movement in healthy and diseased neurons. Frontiers in cellular neuroscience. 2019;13:470.

25. Kumar V, Vasudevan A, Venkatesh K, et al. Cargo crowding, stationary clusters and dynamical reservoirs in axonal transport. bioRxiv. 2021:2021.03. 12.434740.

26. Graf von Keyserlingk D, Schramm U. Diameter of axons and thickness of myelin sheaths of the pyramidal tract fibres in the adult human medullary pyramid. Anatomischer Anzeiger. 1984;157(2):97–111.

27. Verhaart W. On thick and thin fibers in the pyramidal tract. Acta Psychiatrica Scandinavica. 1947;22(3-4):271-281.

28. Fukuda Y, Pazyra-Murphy MF, Silagi ES, et al. Binding and transport of SFPQ-RNA granules by KIF5A/KLC1 motors promotes axon survival. Journal of Cell Biology. 2020;220(1):e202005051.

29. Li W, Sun W, Zhang Y, et al. Rapid induction and long-term self-renewal of primitive neural precursors from human embryonic stem cells by small molecule inhibitors. Proceedings of the National Academy of Sciences. 2011;108(20):8299–8304.

30. Trujillo CA, Gao R, Negraes PD, et al. Complex Oscillatory Waves Emerging from Cortical Organoids Model Early Human Brain Network Development. Cell Stem Cell. Oct 3 2019;25(4):558–569.e7. doi:10.1016/j.stem.2019.08.002

31. Zhao Y, Zhang Y, Teng Y, et al. SpyCLIP: an easy-to-use and high-throughput compatible CLIP platform for the characterization of protein-RNA interactions with high accuracy. Nucleic Acids Res. Apr 8 2019;47(6):e33. doi:10.1093/nar/gkz049

32. Lu L, Liu X, Huang W-K, et al. Robust Hi-C maps of enhancer-promoter interactions reveal the function of non-coding genome in neural development and diseases. Molecular cell. 2020;79(3):521–534. e15.

33. Park Y, Gerson SL. DNA repair defects in stem cell function and aging. Annu Rev Med. 2005;56(1):495–508.

34. Stanganello E, Zahavi EE, Burute M, et al. Wnt signaling directs neuronal polarity and axonal growth. Iscience. 2019;13:318–327.

35. Yi W, Li J, Zhu X, et al. CRISPR-assisted detection of RNA–protein interactions in living cells. Nature methods. 2020;17(7):685–688.

36. Falzone TL, Stokin GB, Lillo C, et al. Axonal stress kinase activation and tau misbehavior induced by kinesin-1 transport defects. Journal of Neuroscience. 2009;29(18):5758–5767.

37. Deng CY, Lei WL, Xu XH, Ju XC, Liu Y, Luo ZG. JIP1 mediates anterograde transport of Rab10 cargos during neuronal polarization. J Neurosci. Jan 29 2014;34(5):1710–23. doi:10.1523/jneurosci.4496-13.2014

38. Kamal A, Stokin GB, Yang Z, Xia C-H, Goldstein LS. Axonal transport of amyloid precursor protein is mediated by direct binding to the kinesin light chain subunit of kinesin-I. Neuron. 2000;28(2):449–459.

39. Sun J, Pan CQ, Chew TW, Liang F, Burmeister M, Low BC. BNIP-H recruits the cholinergic machinery to neurite terminals to promote acetylcholine signaling and neuritogenesis. Developmental Cell. 2015;34(5):555–568.

40. Cosker KE, Fenstermacher SJ, Pazyra-Murphy MF, Elliott HL, Segal RA. The RNA-binding protein SFPQ orchestrates an RNA regulon to promote axon viability. Nature neuroscience. 2016;19(5):690–696.

41. Lim YW, James D, Huang J, Lee M. The emerging role of the RNA-binding protein SFPQ in neuronal function and neurodegeneration. International journal of molecular sciences. 2020;21(19):7151.

42. Takamori S, Holt M, Stenius K, et al. Molecular anatomy of a trafficking organelle. Cell. 2006;127(4):831–846.

43. Misgeld T, Kerschensteiner M, Bareyre FM, Burgess RW, Lichtman JW. Imaging axonal transport of mitochondria in vivo. Nature methods. 2007;4(7):559–561.

44. Alonso AdC, Grundke-Iqbal I, Iqbal K. Alzheimer’s disease hyperphosphorylated tau sequesters normal tau into tangles of filaments and disassembles microtubules. Nature medicine. 1996;2(7):783–787.

45. Elvira G, Wasiak S, Blandford V, et al. Characterization of an RNA granule from developing brain. Molecular & cellular proteomics. 2006;5(4):635–651.

46. Levone BR, Lenzken SC, Antonaci M, et al. FUS-dependent liquid–liquid phase separation is important for DNA repair initiation. Journal of Cell Biology. 2021;220(5):e202008030.

47. Xiao M, Wang F, Chen N, et al. Smad4 sequestered in SFPQ condensates prevents TGF-β tumor-suppressive signaling. Developmental Cell. 2024;59(1):48–63. e8.

48. Zhao Y, Zhang Y, Teng Y, et al. SpyCLIP: an easy-to-use and high-throughput compatible CLIP platform for the characterization of protein–RNA interactions with high accuracy. Nucleic acids research. 2019;47(6):e33–e33.

49. Tichon A, Gil N, Lubelsky Y, et al. A conserved abundant cytoplasmic long noncoding RNA modulates repression by Pumilio proteins in human cells. Nature communications. 2016;7(1):12209.

50. Hirokawa N, Niwa S, Tanaka Y. Molecular motors in neurons: transport mechanisms and roles in brain function, development, and disease. Neuron. 2010;68(4):610–638.

51. Konecna A, Frischknecht R, Kinter J, et al. Calsyntenin-1 docks vesicular cargo to kinesin-1. Molecular biology of the cell. 2006;17(8):3651–3663.

52. Dimitrova-Paternoga L, Jagtap PKA, Cyrklaff A, et al. Molecular basis of mRNA transport by a kinesin-1–atypical tropomyosin complex. Genes & development. 2021;35(13-14):976–991.

53. Hirokawa N. mRNA transport in dendrites: RNA granules, motors, and tracks. Journal of Neuroscience. 2006;26(27):7139–7142.

54. McNally KL, Martin JL, Ellefson M, McNally FJ. Kinesin-dependent transport results in polarized migration of the nucleus in oocytes and inward movement of yolk granules in meiotic embryos. Developmental biology. 2010;339(1):126–140.

55. Pichon X, Moissoglu K, Coleno E, et al. The kinesin KIF1C transports APC-dependent mRNAs to cell protrusions. Rna. 2021;27(12):1528–1544.

56. Ganesh VS, Riquin K, Chatron N, et al. Neurodevelopmental Disorder Caused by Deletion of CHASERR, a lncRNA Gene. New England Journal of Medicine. 2024;391(16):1511–1518.

57. Ramos AD, Andersen RE, Liu SJ, et al. The long noncoding RNA Pnky regulates neuronal differentiation of embryonic and postnatal neural stem cells. Cell stem cell. 2015;16(4):439–447.

58. Zhao X, Tang Z, Zhang H, et al. A long noncoding RNA contributes to neuropathic pain by silencing Kcna2 in primary afferent neurons. Nature neuroscience. 2013;16(8):1024–1031.

59. Sauvageau M, Goff LA, Lodato S, et al. Multiple knockout mouse models reveal lincRNAs are required for life and brain development. elife. 2013;2:e01749.

60. Chalei V, Sansom SN, Kong L, et al. The long non-coding RNA Dali is an epigenetic regulator of neural differentiation. elife. 2014;3:e04530.

61. Vance KW, Sansom SN, Lee S, et al. The long non-coding RNA P aupar regulates the expression of both local and distal genes. The EMBO journal. 2014;33(4):296–311.

62. Cajigas I, Leib DE, Cochrane J, et al. Evf2 lncRNA/BRG1/DLX1 interactions reveal RNA-dependent inhibition of chromatin remodeling. Development. 2015;142(15):2641–2652.

63. Raveendra BL, Swarnkar S, Avchalumov Y, et al. Long noncoding RNA GM12371 acts as a transcriptional regulator of synapse function. Proceedings of the National Academy of Sciences. 2018;115(43):E10197–E10205.

64. Lin MF, Jungreis I, Kellis M. PhyloCSF: a comparative genomics method to distinguish protein coding and non-coding regions. Bioinformatics. 2011;27(13):i275–282.

65. Kopp F, Elguindy MM, Yalvac ME, et al. PUMILIO hyperactivity drives premature aging of Norad-deficient mice. Elife. 2019;8:8:e42650.

66. Sasaki E, Suemizu H, Shimada A, et al. Generation of transgenic non-human primates with germline transmission. Nature. 2009;459(7246):523–527.

67. Courchesne E, Mouton PR, Calhoun ME, et al. Neuron number and size in prefrontal cortex of children with autism. The Journal of the American Medical Association. 2011;306(18):2001–2010.

68. Yoon SI, Kurnasov O, *Natarajan* V, et al. Structural basis of TLR5-flagellin recognition and signaling. Science. 2012;335(6070):859–864.

69. Koser DE, Thompson AJ, Foster SK, et al. Mechanosensing is critical for axon growth in the developing brain. Nature neuroscience. 2016;19(12):1592–1598.

70. Saez-Atienzar S, Masliah E. Cellular senescence and Alzheimer disease: the egg and the chicken scenario. Nature reviews neuroscience. 2020;21(8):433–444.

71. Thomas-Jinu S, Gordon PM, Fielding T, et al. Non-nuclear Pool of Splicing Factor SFPQ Regulates Axonal Transcripts Required for Normal Motor Development. Neuron. 2017;94(2):322–336.

